# *Bartonella taylorii*: A model organism for studying *Bartonella* infection *in vitro* and *in vivo*

**DOI:** 10.1101/2022.03.11.483615

**Authors:** Katja Fromm, Alexandra Boegli, Monica Ortelli, Alexander Wagner, Erwin Bohn, Silke Malmsheimer, Samuel Wagner, Christoph Dehio

## Abstract

*Bartonella* spp. are Gram-negative facultative intracellular pathogens that infect diverse mammals and cause a long-lasting intra-erythrocytic bacteremia in their natural host. These bacteria translocate *Bartonella* effector proteins (Beps) into host cells via their VirB/VirD4 type 4 secretion system (T4SS) in order to subvert host cellular functions, thereby leading to the downregulation of innate immune responses. Most studies on the functional analysis of the VirB/VirD4 T4SS and the Beps were performed with the major zoonotic pathogen *Bartonella henselae* for which efficient *in vitro* infection protocols have been established. However, its natural host, the cat, is unsuitable as an experimental infection model. *In vivo* studies were mostly confined to rodent models using rodent-specific *Bartonella* species, while the *in vitro* infection protocols devised for *B. henselae* are not transferable for those pathogens. The disparities of *in vitro* and *in vivo* studies in different species has hampered progress in our understanding of *Bartonella* pathogenesis. Here we describe the murine-specific strain *B. taylorii* IBS296 as a new model organism facilitating the study of bacterial pathogenesis both *in vitro* in cell cultures and *in vivo* in laboratory mice. We implemented the split NanoLuc luciferase-based translocation assay to study BepD translocation through the VirB/VirD4 T4SS. We found increased effector-translocation into host cells if the bacteria were grown on tryptic soy agar (TSA) plates and experienced a temperature shift immediately before infection. The improved infectivity *in vitro* could be correlated to an upregulation of the VirB/VirD4 T4SS. Using our adapted infection protocols, we showed BepD-dependent immunomodulatory phenotypes *in vitro.* In mice, the implemented growth conditions enabled infection by a massively reduced inoculum without having an impact on the course of the intra-erythrocytic bacteremia. The established model opens new avenues to study the role of the VirB/VirD4 T4SS and the translocated Bep effectors *in vitro* and *in vivo.*

## 1 Introduction

Bartonellae are Gram-negative facultative intracellular pathogens, which infect diverse mammals including humans. Clinically relevant infections with *Bartonella* are caused by zoonotic *Bartonella henselae*, the agent of the cat scratch disease (CSD) (Huarcaya et al., 2002; Khalfe and Lin, 2022) or human-specific species such as *Bartonella bacilliformis*, the agent of the life-threatening Carrion’s disease, and *Bartonella quintana*, which causes trench fever (Maguina et al., 2009; Mada et al., 2022). Bartonellae are highly host-restricted pathogens. After transmission by an arthropod vector, the bacteria enter the dermis and eventually seed into the blood stream where they cause a long-lasting intra-erythrocytic bacteremia as hallmark of infection in their natural host (Seubert et al., 2002; Chomel et al., 2009; Harms and Dehio, 2012). Infections of incidental hosts are not associated with intra-erythrocytic persistence but clinical manifestations caused by several zoonotic species can range from mild symptoms to severe diseases (Chomel and Kasten, 2010; Wagner and Dehio, 2019).

Previous studies demonstrated that the progression to the blood stream requires a functional VirB/VirD4 type 4 secretion system (T4SS) (Schulein and Dehio, 2002). T4SS are multi-protein complexes embedded into the cell envelop. In Bartonellae, VirB2-11 are assembled to the functional T4SS that facilitates substrate translocation. The ATPase VirD4, also referred to as the type 4 secretion (T4S) coupling protein (T4CP), is essential for substrate recognition and entry into the T4SS (Berge et al., 2017; Waksman, 2019). In *Bartonella* species the *virB2* promoter *(PvirB2)* drives expression of the *virB2-11 operon.* This promoter and the separate promoter of *virD4* are controlled by the BatR/BatS two-component system. Upregulated expression of the VirB/VirD4 T4SS in *B. henselae* is linked to the BatR/BatS two-component system activated at physiological pH and the alternative sigma factor RpoH1. Induction of RpoH1 is mediated by the stringent response, which relies on the accumulation of the second messenger guanosine tetra- and pentaphosphate (both referred to as ppGpp) in the bacterial cytosol. (Quebatte et al., 2010; Quebatte et al., 2013).

The VirB/VirD4 T4SS is important to translocate multiple *Bartonella* effector proteins (Beps) into mammalian host cells to subvert host cellular functions, e.g. to dampen innate immune responses (Wagner et al., 2019; Fromm and Dehio, 2021). Different *Bartonella* species translocate discrete cocktails of Beps into host cells (Harms et al., 2017b). While some orthologs share a conserved function (Sorg et al., 2020), other Beps seem to vary in a species-specific manner (Schmid et al., 2006a; Wang et al., 2019). Extensive studies focusing on the role of the VirB/VirD4 T4SS and the translocated Beps have been performed *in vitro* and *in vivo*. However, experimental studies on these bacteria in the natural host share the problem that either the host as model is hardly available or protocols for *in vitro* studies are missing. *B. henselae* is among the best-characterized *Bartonella* species and *in vitro* infection protocols using various cell lines or primary cells were published (Musso et al., 2001; McCord et al., 2005; Ma and Ma, 2016; Sorg et al., 2020; Marlaire and Dehio, 2021). Investigating *B. henselae* in its natural host, the cat, is laborious and expensive (Chomel et al., 1996; Foil et al., 1998). In a mouse infection model *B. henselae* failed to establish long-lasting intra-erythrocytic bacteremia and pathology also differed from infections in the natural host (Regnath et al., 1998; Kunz et al., 2008). On the other hand, several rodent infection models with rodent-specific species were published that recapitulate the long-lasting intra-erythrocytic infection course characteristic for the natural host (Boulouis et al., 2001; Koesling et al., 2001; Schulein and Dehio, 2002; Deng et al., 2016; Siewert et al., 2021). However, besides an erythrocyte invasion model no *in vitro* protocols were established to study the interaction of those species with cells of their natural hosts (Vayssier-Taussat et al., 2010). Establishing for at least one *Bartonella* strain an experimental model for *in vitro* and *in vivo* studies would help studying the role of the VirB/VirD4 T4SS and its translocated Beps in the context of infection in the natural reservoir.

In this study, we investigated *Bartonella taylorii* IBS296 as a model organism to study VirB/VirD4 and Bep effector-related functions *in vitro* and *in vivo.* We confirmed that *B. taylorii* is dampening the innate immune response *in vitro* in a VirB/VirD4 and BepD-dependent manner. Further, we established the split NanoLuc luciferase-based translocation assay (Westerhausen et al., 2020) to study BepD translocation through the VirB/VirD4 T4SS. In addition, we improved the previously established mouse infection model for *B. taylorii* (Siewert et al., 2021) by lowering the inoculum by several orders of magnitude without affecting course of bacteremia. Our findings provide the basis for more extensive studies focusing on the immunomodulatory function of *B. taylorii in vitro* and *in vivo*.

## 2 Materials and methods

### 2.1 Bacterial Strains and growth conditions

All bacterial strains used in this study are listed in table 1. *E. coli* strains were cultivated in lysogeny broth (LB) or on solid agar plates (LA) supplemented with appropriate antibiotics at 37°C overnight.

**Table 1.** List and construction of all bacterial strains of this study

| Strain | Genotype | Reference/Source | Identifier/Description |
| --- | --- | --- | --- |
| ***Escherichia coli*** |  |  |  |
| Novablue | *endA1 hsdR17 (r<sub>K12</sub><sup>-</sup> m<sub>K12</sub><sup>+</sup>) supE44 thi-1 recA1 gyrA96 relA1 lac F'[proA<sup>+</sup>B<sup>+</sup> lacI<sup>q</sup>ZΔM15::Tn10]* | Novagen | Standard cloning strain |
| HST08 | *F<sup>-</sup>, endA1, supE44, thi-1, recA1, relA1, gyrA96, phoA, Φ80d lacZΔM15, Δ (lacZYA - argF) U169, Δ (mrr - hsdRMS - mcrBC), ΔmcrA, λ-* | Takara | Standard cloning strain |
| JKE201 | MFDpir Δ*mcrA* Δ(*mrr-hsdRMS-mcrBC*) *aac(3)IV::lacI<sup>q</sup>* | (Harms et al., 2017b) | derivative of MFDpir lacking EcoKI, the three type IV restriction systems, restored gentamicin sensitivity, harboring *lacI<sup>q</sup>* allele |
| ***Bartonella henselae*** |  |  |  |
| *B. henselae* Houston-1 | *rpsL* | (Schmid et al., 2004) | RSE247, spontaneous SmR strain of *B. henselae* ATCC49882T, serving as wild-type |
|  | *rpsL pCD366* | (Quebatte et al., 2010) | MQB1610; RSE247 containing pCD366 |
|  | *rpsL pAH196* | (Harms et al., 2017b) | MQB1612; RSE247 containing pAH196_Bhe |
|  | *rpsL ΔvirD4* | (Schulein et al., 2005) | GS0221; virD4 deletion mutant, derivative of RSE247 |
|  | *rpsL ΔvirD4 / pHiBiT-FLAG* | This study | KFB286; GS0221 containing pKF059 |
|  | *rpsL ΔvirD4 / pHiBiT-FLAG-bepD<sub>Bhe</sub>* | This study | MOB120; GS0221 containing pMO006 |
|  | *rpsL ΔbepA-G* | (Schulein et al., 2005) | MSE150; bepA-bepG deletion mutant, derivative of RSE247 |
|  | *rpsL ΔbepA-G / pbepD<sub>Bhe</sub>* | (Schulein et al., 2005) | PG4D03; MSE150 containing pPG104 |
|  | *rpsL ΔbepA-G / pbepD<sub>Bta</sub>* | (Sorg et al., 2020) | LUB242; MSE150 containing pLU058 |
|  | *rpsL ΔbepA-G / pHiBiT-FLAG* | This study | KFB276; MSE150 containing pKF059 |
|  | *rpsL ΔbepA-G / pHiBiT-FLAG-bepD<sub>Bhe</sub>* | This study | MOB121; MSE150 containing pMO006 |
| ***Bartonella***<br>***taylorii*** |  |  |  |
| *B. taylorii*<br>IBS296 Sm<sup>R</sup> | *rpsL* | (Sorg et al., 2020) | KFB030, spontaneous SmR strain of *B. taylorii* IBS296, serving as wild-type, derivative of LUB046 |
|  | *rpsL pCD366* | This study | KFB266; KFB030 containing pCD366 |
|  | *rpsL pAH196_Btay* | (Harms et al., 2017b) | KFB063; LUB046 containing pAH196_Btay |
|  | *rpsL ΔvirD4* | This study | KFB146; *virD4* deletion mutant of KFB030 |
|  | *rpsL ΔvirD4 / pHiBiT-FLAG* | This study | KFB291, KFB146 containing pKF059 |
|  | *rpsL ΔvirD4 / pHiBiT-FLAG-bepD<sub>Bta</sub>* | This study | KFB233, KFB146 containing pKF027 |
|  | *rpsL ΔbepA* | This study | KFB068; *bepA* deletion mutant of KFB030 |
|  | *rpsL ΔbepC* | This study | KFB085, *bepC* deletion mutant of KFB030 |
|  | *rpsL ΔbepD* | This study | KFB070; *bepD* deletion mutant of KFB030 |
|  | *rpsL ΔbepF* | This study | KFB097; *bepF* deletion mutant of KFB030 |
|  | *rpsL ΔbepI* | This study | KFB101; *bepI* deletion mutant of KFB030 |
|  | *rpsL ΔbepA-I* | This study | KFB072; *bepA-bepI* deletion mutant of KFB030 |
|  | *rpsL ΔbepA-I / pHiBiT-FLAG* | This study | KFB287; KFB072 containing pKF059 |
|  | *rpsL ΔbepA-I / pHiBiT-FLAG-bepD<sub>Bta</sub>* | This study | KFB263; KFB072 containing pKF027 |

*Bartonella* strains were grown at 35°C and 5% CO_2_ on Columbia blood agar (CBA) or tryptic soy agar (TSA) plates supplemented with 5% defibrinated sheep blood and appropriate antibiotics. *Bartonella* strains stored as frozen stocks at −80°C were inoculated on CBA or TSA plates for 3 days and subsequently expanded on fresh plates for 2 days. Prior to infection *Bartonella* strains were cultured in M199 medium supplemented with 10% heat-inactivated fetal calf serum (FCS) for 24 h at an optical density (OD_600 nm_) of 0.5 at 28°C or 35°C and 5% CO_2_.

Antibiotics or supplements were used in the following concentrations: kanamycin at 30 μg/ml, streptomycin at 100 μg/ml, isopropyl-b-D-thiogalactoside (IPTG) at 100 μM and diaminopimelic acid (DAP) at 1 mM.

### 2.2 Construction of strains and plasmids

DNA manipulations were performed according to standard techniques and all cloned inserts were DNA sequenced to confirm sequence integrity. For protein complementation/overexpression in *Bartonella* selected genes were cloned into plasmid pBZ485_empty under the control of the taclac promoter. Chromosomal deletions or insertions of *B. taylorii* were generated by a two-step gene replacement procedure as previously described (Schulein and Dehio, 2002). A detailed description for the construction of each plasmid is presented in table 2. The sequence of all oligonucleotide primers used in this study is listed in table 3.

**Table 2:** List of plasmids used in this study

| Plasmid | Backbone | Description | Reference/Source |
| --- | --- | --- | --- |
| pCD366 |  | RSF1010 derivative encoding promoterless gfpmut2 | (Dehio et al., 1998) |
| pAH196_Bhe | pCD366 | pCD366 with PvirB2 of *B. henselae* ahead of gfpmut2 | (Harms et al., 2017b) |
| pAH196_Btay | pCD366 | pCD366 with PvirB2 of *B. taylorii* ahead of gfpmut2 | (Harms et al., 2017b) |
| pTR1000 |  | *Bartonella* suicide plasmid with *kanR* / *rpsL* double-selectable cassette for scarless deletions | (Schulein and Dehio, 2002) |
| pKF001 | pTR1000 | pTR1000 with homology sites to delete *bepA* of *Bartonella taylorii* (homology regions amplified separately and then fused by SOEing PCR) | This study |
| pKF002 | pTR1000 | pTR1000 with homology sites to delete *bepD* of *Bartonella taylorii* (homology regions amplified separately and then fused by SOEing PCR) | This study |
| pKF003 | pTR1000 | pTR1000 with homology sites to delete *bepC-I* of *Bartonella taylorii* (homology regions amplified separately and then fused by SOEing PCR) | This study |
| pKF005 | pTR1000 | pTR1000 with homology sites to delete *bepC* of *Bartonella taylorii* (homology regions amplified separately and then fused by SOEing PCR) | This study |
| pKF006 | pTR1000 | pTR1000 with homology sites to delete *bepF* of *Bartonella taylorii* (homology regions amplified separately and then fused by SOEing PCR) | This study |
| pKF007 | pTR1000 | pTR1000 with homology sites to delete *bepI* of *Bartonella taylorii* (homology regions amplified separately and then fused by SOEing PCR) | This study |
| pKF008 | pTR1000 | pTR1000 with homology sites to delete *virD4* of *Bartonella taylorii* (homology regions amplified separately and then fused by SOEing PCR) | This study |
| pBZ485 |  | new *E. coli* / *Bartonella* shuttle vector based on pCD341 with *Plac*(MQ5); RP4 *oriT* | (Harms et al., 2017a) |
| pKF059 | pBZ485 | Derivative of pBZ485, encodes for HiBiT::FLAG | This study |
| pKF027 | pBZ485 | Derivative of pBZ485, encodes for HiBiT::FLAG *Bta* BepD fusion protein | This study |
| pMO006 | pBZ485 | Derivative of pBZ485, encodes for HiBiT::FLAG *Bhe* BepD fusion protein | This study |

**Table 3:**
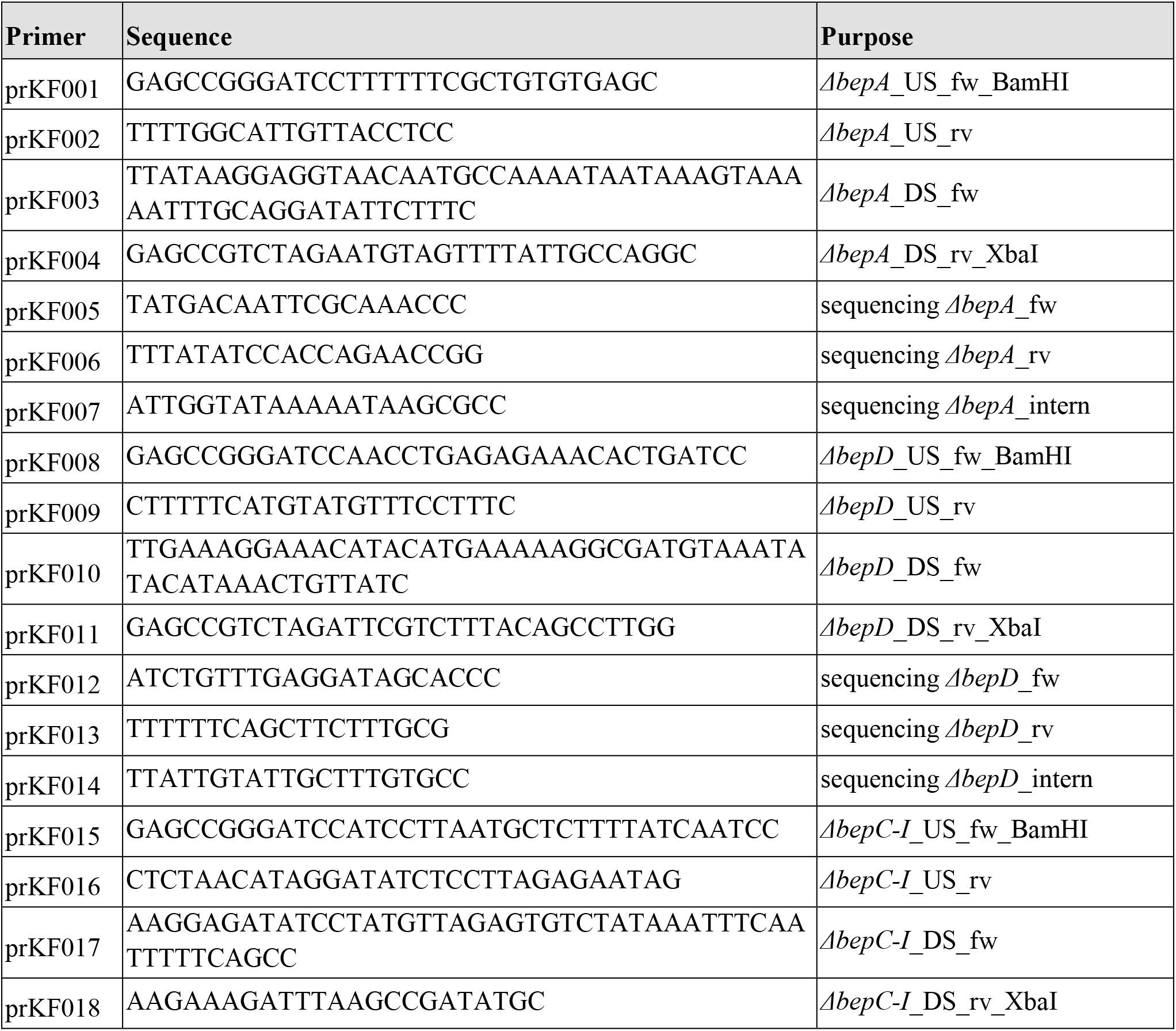
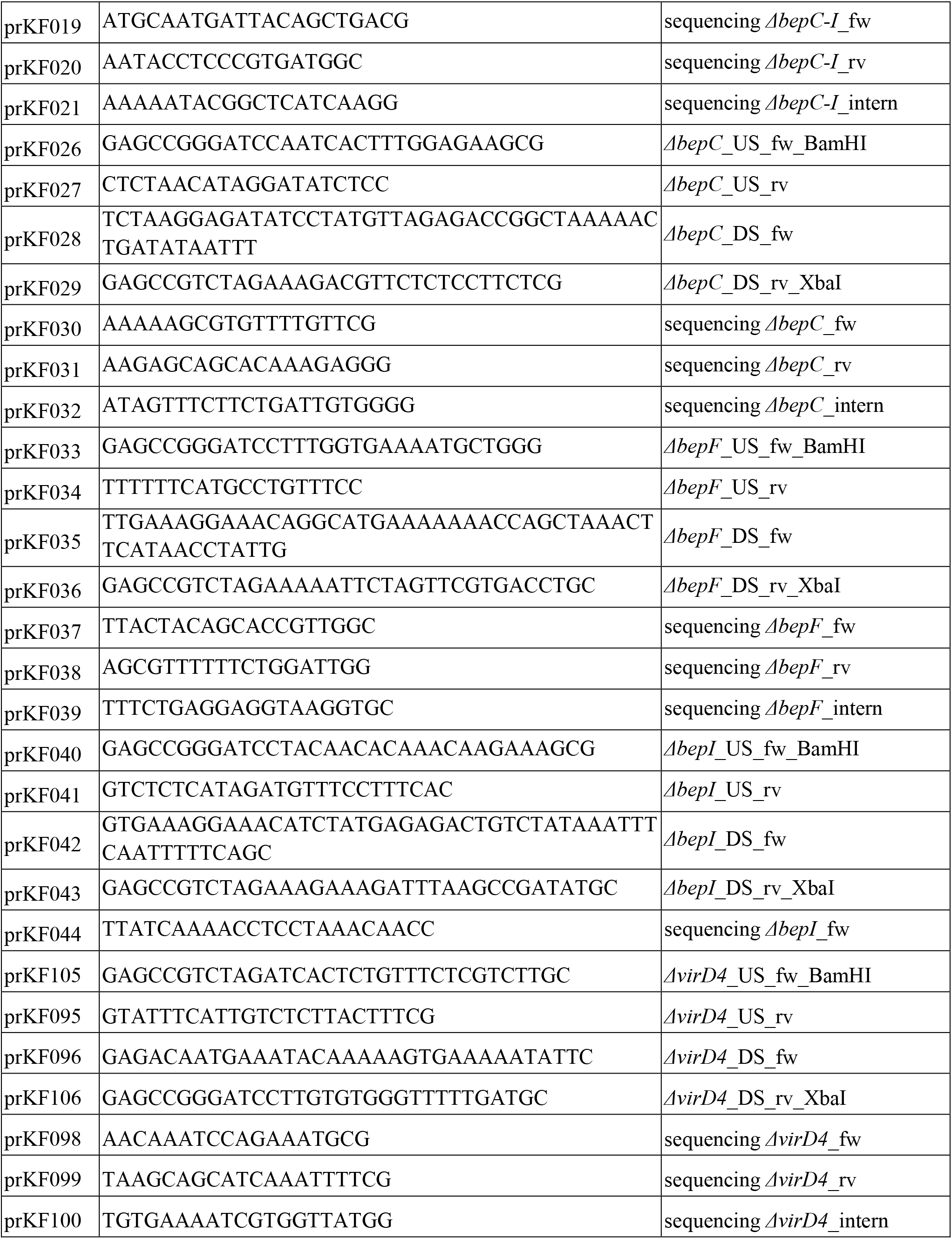
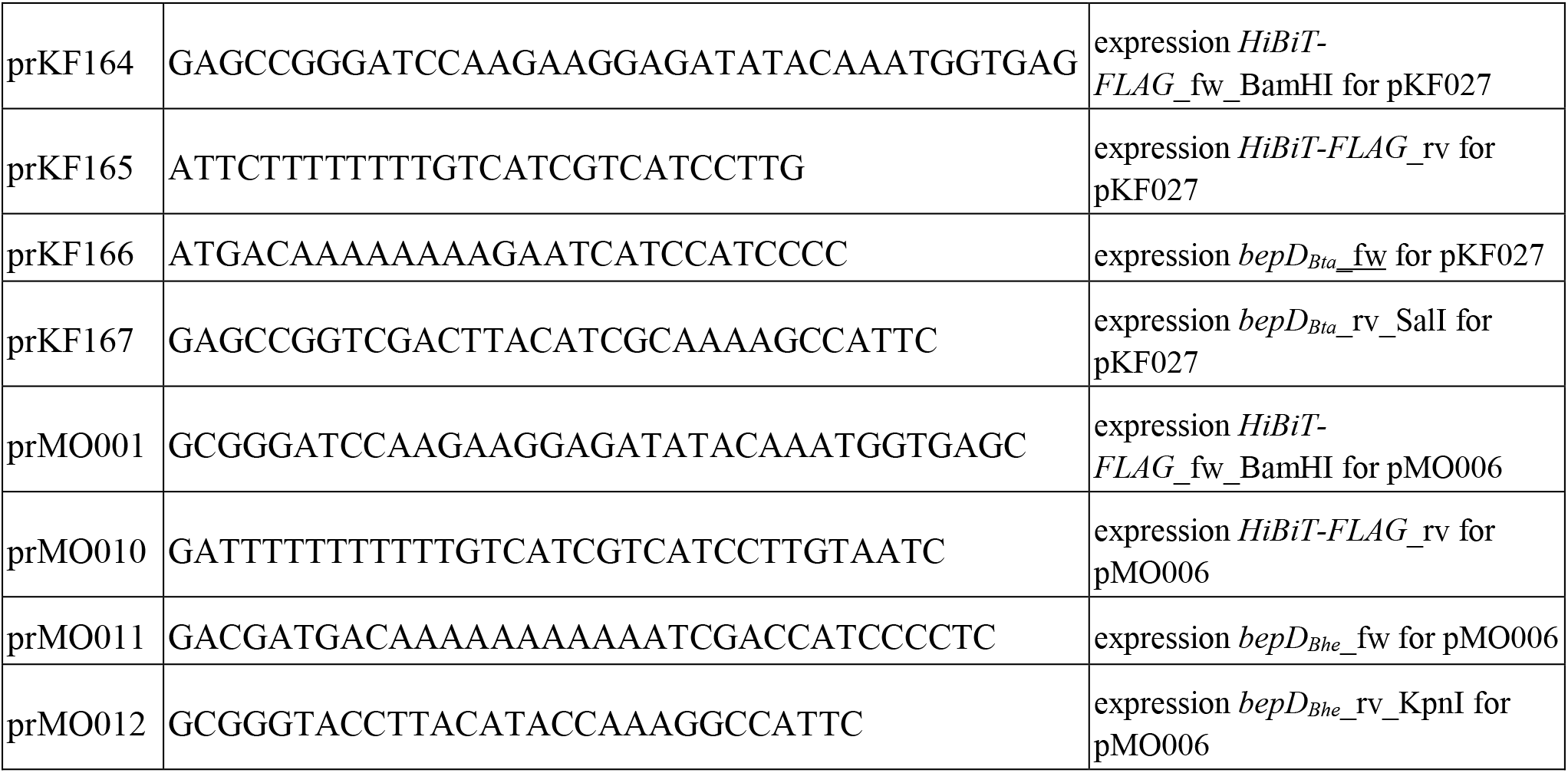
List of oligonucleotide primers used in this study

Plasmids were introduced into *Bartonella* strains by conjugation from *E. coli* strain MFDpir using two-parental mating as described previously (Harms et al., 2017b).

### 2.3 Cell Lines and Culture Conditions

JAWS II (ATCC CRL-11904) cell line is a GM-CSF-dependent DC line established from bone marrow cells of a p53-knockout C57BL/6 mouse (Jiang et al., 2008). JAWS II cells were cultured at 37°C in 5% CO_2_ in complete culture medium consisting of MDM with 20% FCS, 4 mM L-glutamine, 1 mM sodium pyruvate and 5 ng/ml GM-CSF. RAW 264.7 (ATCC TIB-71) cell line is a murine macrophage cell line originating from an adult male BALB/c mouse (Raschke et al., 1978). RAW 264.7 and RAW 264.7 LgBiT (obtained from S. Wagner, University Tübingen, Germany) cells were cultured at 37°C and 5% CO_2_ in DMEM Glutamax supplemented with 10% FCS. RAW 264.7 LgBiT cells were treated with 2 ng/mL puromycin to select for stably transduced cells.

### 2.4 Cell Infections

*B. henselae* and *B. taylorii* strains were cultured as described above. One day before infection, 1 x 10^5^ cells (JAWS II) or 5 x 10^5^ cells (RAWs) were seeded per well in 12-well plates. 1 x 10^4^ cells (RAWs LgBiT) per well were seeded in white 96-well plates (Corning, no. CLS3610). 1 h prior to infection bacterial cultures were supplemented with 100 μM IPTG to induce protein expression, if required. Cells were washed once with infection medium (DMEM Glutamax, supplemented with 1% FCS) and infected with a multiplicity of infection (MOI) of 50 bacteria per cell, if not stated otherwise. Bacterial attachment was synchronized by centrifugation at 500 g for 3 min. Infected cells were incubated at 37°C and 5% CO_2_ for the indicated times. If indicated, cells were stimulated with 100 ng/ml LPS (lipopolysaccharides from *E. coli* O26:B6, Sigma-Aldrich) during the last 2 h of infection at 37°C and 5% CO_2_. Supernatants were analyzed by Ready-SET-Go! ELISA kits for TNF-α. Adherent cells were harvested, lysed and analyzed by immunoblot. For monitoring effector injection, luminescence reading was carried out in the Synergy H4 plate reader (BioTek).

### 2.5 Quantification of Cytokine Levels in Culture Supernatants

TNF-α was quantified in cell culture supernatants of infected cells by Ready-SET-Go! ELISA kits (Thermo Fisher Scientific, Cat. 88-7324-77) according to the manufacturer’s instructions. 96-well assay plates (Costar no. 9018) were coated overnight at 4°C with capture antibody in coating buffer. The plates were washed with wash buffer (PBS containing 0.05% Tween-20) and incubated at RT for 1 h with assay diluent (provided in the kit) to block unspecific binding. Samples were added directly to the plate or after pre-dilution in assay diluent. Pre-dilutions used for TNF-α quantification: JAWS II (1 to 4 for samples with LPS stimulation; no pre-dilution for unstimulated samples), RAW264.7 (1 to 6for samples with LPS stimulation; no pre-dilution for unstimulated samples). Subsequently samples were diluted twice by serial 2-fold dilutions on the plate. The respective lyophilized standard was resolved as requested, added to the plate and diluted by serial 2-fold dilutions. The plate was incubated at 4°C overnight. 5 wash steps were performed. After addition of the respective detection antibody, the plate was incubated 1 hour at RT. Horseradish peroxidase-conjugated avidin was added for 30 min at room temperature after another 5 washing steps. After 7 washes ELISA substrate solution was added for 5 to 15 min at RT and the reaction stopped by adding 1M H3PO4. Absorbance was read at 450 and 570 nm. At least three independent experiments (n = 3) were performed in technical triplicates.

### 2.6 SDS-PAGE, western blotting and immunodetection

SDS-PAGE and immunoblotting were performed as described (Schulein et al., 2005). To verify expression levels of the protein of interest, JAWS II or RAW 264.7 cells were collected and washed in ice-cold PBS. Cell pellets were lysed by adding Novagen‘s PhosphoSafe extraction buffer (Merck, Cat. 71296) complemented with cOmplete Mini EDTA-free protease inhibitor cocktail (Roche, 11836170001). Lysate protein concentrations were quantified using the Pierce BCA Protein Assay kit (Thermo Fisher Scientific, Cat. 23225). Lysates with equal protein concentrations were mixed with 5x SDS sample buffer, and resolved on 4 – 20% precast protein TGX gels (BioRad, Cat. 456-1093, Cat. 456-1096). Pre-stained Precision Plus Protein Dual Color Standard (BioRad, Cat. 1610374) was used as protein size reference. Proteins were transferred onto Amersham Protran® Nitocellulose Blotting membrane (0.45 μm pore size) or Amersham Hybond® PVDF membrane (0.2 μm pore size). Membranes were probed with primary antibodies directed against the protein of interest (α-actin (Milipore, Cat. MAB1501), α-pSTAT3 (Y705) (Cell Signaling Technology, Cat. 9145), α-STAT3 (Cell Signaling Technology, Cat. 12640), α-FLAG (Sigma-Aldrich, Cat. F1804). Detection was performed with horseradish peroxidase-conjugated antibodies directed against rabbit or mouse IgG (HRP-linked α-mouse IgG (Cell Signaling Technology, Cat. 7076), HRP-linked α-rabbit IgG (Cell Signaling Technology, Cat. 7074)). Immunoblots were developed using LumiGLO® chemiluminescent substrate (Seracare, Cat. 5430) and imaged using the Fusion FX device (Vilber). Signal quantification was performed using the Fusion FX7 Edge software. If required, images were adjusted in brightness and contrast using the ImageJ software.

### 2.7 Determination of promoter expression by flow cytometry

Bartonellae were grown on CBA or TSA plates as described above. Bacteria were resuspended in M199 + 10% FCS, diluted to a final concentration of OD_600 nm_ = 0.5 and incubated for 24 h at 28°C or 35°C at 5% CO_2_. The bacteria were harvested, centrifuged at 1900 g for 4 min and washed in FACS buffer (2% FCS in PBS). After another centrifugation step, the supernatant was aspirated, bacteria fixed in 3.7% PFA for 10 min at 4°C and finally resuspended in FACS buffer. Expression of the *PvirB2:gfp* promoter was evaluated by measuring the GFP fluorescence signal using the BD LSRFortessa. Data analysis was performed using FlowJo v10.6.2.

### 2.8 NanoLuc-based effector translocation assay

To assess whether the HiBiT-FLAG fragment, HiBiT-FLAG-BepD_*Bhe*_ and HiBiT-FLAG-BepD_*Bta*_ can complement LgBiT to a functional luciferase, we used the Nano-Glo HiBiT lytic detection system (Promega, Cat. N3030). Bacteria were cultured as previously indicated. 5 x 10^8^ bacteria were resuspended in 100 μL PBS and supplemented with 100 μL LSC buffer containing the substrate (1/50 v/v) and the LgBiT protein (1:100 v/v). The luminescent signal was measured using the Synergy H4 plate reader.

RAW LgBiT macrophages were infected as described previously. Effector translocation into macrophages was quantified by measuring the luminescent signal using the Synergy H4 plate reader. The Nano-Glo live cell reagent (Promega, Cat. N2011) was prepared as advised by manufacturer. After 24 h of infection, supernatant was aspirated and cells were gently washed in pre-warmed PBS. A final volume of 100 uL PBS per well was supplemented with 25 uL of the Nano-Glo live cell assay buffer containing the substrate for luminescence measurement in the Synergy H4 plate reader. The following settings were used: temperature 37°C, shaking sequence 30 sec at 300-500 rpm, delay 10 min, autoscale, integration time 5 sec. At least three independent experiments (n = 3) were performed in technical triplicates.

### 2.9 Mouse experimentation

Mice handling was performed in strict accordance with the Federal Veterinary Office of Switzerland and local animal welfare bodies. All animal work was approved by the Veterinary Office of the Canton Basel City (license number 1741). All animals were housed at SPF (specific pathogen free) conditions and provided water and food ad libitum. The animal room was on a 12 light/12 dark cycle, and cage bedding changed every week. Female C57BL/6JRj mice were purchased from Janvier Labs.

Animals were infected *i.d.* with the indicated cfu bacteria in PBS. Blood was drawn in 3.8 % sodium citrate from the tail vein several days post infection. Whole blood was frozen at −80°C, thawed and plated on CBA plates in serial dilutions to determine blood cfu count.

### 2.10 Statistical Analysis

Graphs were generated with GraphPad Prism 8. Statistical analyses were performed using one-way ANOVA with multiple comparisons (Tukey’s multiple comparison test). For the graphs presented in the figures, significance was denoted as non-significant (ns) (p > 0.05); * p < 0.05; ** p < 0.01; *** p < 0.001; **** p < 0.0001. Number of independent biological replicates is indicated as n in the figure legends.

## 3 Results

### 3.1 Establishment of a cell culture assay for BepD translocation via the *B. taylorii* VirB/VirD4 T4SS

*In vitro* infection protocols devised for *B. henselae* enable an efficient and fast translocation of Beps via the VirB/VirD4 T4SS into eukaryotic host cells (Schmid et al., 2006b; Sorg et al., 2020). Inside host cells, the *Bartonella* effector protein D of *B. henselae* (BepD_*Bhe*_) interacts via phospho-tyrosine domains (pY domains) with the transcription factor Signal Transducer and Activator of Transcription 3 (STAT3) and the Abelson tyrosine kinase (c-ABL). c-ABL than phosphorylates STAT3 on Y705 resulting in a downregulation of pro-inflammatory cytokine secretion. The orthologue effector encoded in *B. taylorii, BepD_Bta_*, is also translocated and exhibits a conserved function when ectopically expressed in *B. henselae* (Sorg et al., 2020). We ectopically expressed BepD_*Bhe*_ and BepD_*Bta*_ in *B. henselae ΔbepA-G*, a genomic deletion mutant lacking all *bep* genes. We infected the mouse dendritic cell line JAWS II at a multiplicity of infection (MOI) of 50 for 6 h. Bacteria expressing BepD_*Bhe*_ or BepD_*Bta*_ facilitated STAT3 phosphorylation to a similar extend ((Sorg et al., 2020), suppl. Figure S1A). To establish an *in vitro* infection protocol for *B. taylorii*, which enables the effector translocation via the VirB/VirD4 T4SS, we used the BepD-dependent STAT3 phosphorylation as sensitive readout.

We infected JAWS II cells with *B. taylorii* IBS296 Sm^R^, used as wild-type strain, and the T4S-deficient *ΔvirD4* mutant. Corresponding *B. henselae* strains served as controls. The bacteria were grown on Columbia blood agar (CBA) plates and the VirB/VirD4 system was induced by overnight culturing in M199 + 10% FCS (Quebatte et al., 2013; Sorg et al., 2020). We performed time-course experiments to compare the STAT3 phosphorylation at early time-points after infection. We quantified the phosphorylated STAT3 over the total STAT3. Cells infected for 24 h with the translocation-deficient *ΔvirD4* mutants served as controls and did not show enhanced STAT3 activation. *B. henselae* wild-type triggered STAT3 phosphorylation 1 hour post infection (hpi). However, *B. taylorii* did not induce STAT3 phosphorylation in JAWS II cells within 6 hpi (Figure 1A and 1B). STAT3 phosphorylation was observed only after 24 h in cells infected with *B. taylorii* (suppl. Figure S1B). Since BepD of *B. taylorii* triggered the STAT3 phosphorylation at early time points when expressed in *B. henselae* (suppl. Figure S1A), we speculated that other than described for the *B. henselae* VirB/VirD4 T4SS, the one of *B. taylorii* was not induced during the first hours of infection. To test whether the induction of VirB/VirD4 T4SS can be further enhanced, we optimized the culture conditions. In previous studies, different *Bartonella* species isolated from their natural hosts were cultured on tryptic soy agar (TSA) plates instead of CBA (Li et al., 2015; Stepanic et al., 2019). Compared to the *ΔvirD4* mutant, *B. taylorii* wild-type harvested from TSA plates induced STAT3 phosphorylation already after 3 hpi. At 6 hpi cells infected with *B. taylorii* showed STAT3 phosphorylation to comparable levels as triggered by *B. henselae* grown on CBA (Figure 1C and 1D).

**Figure 1:**
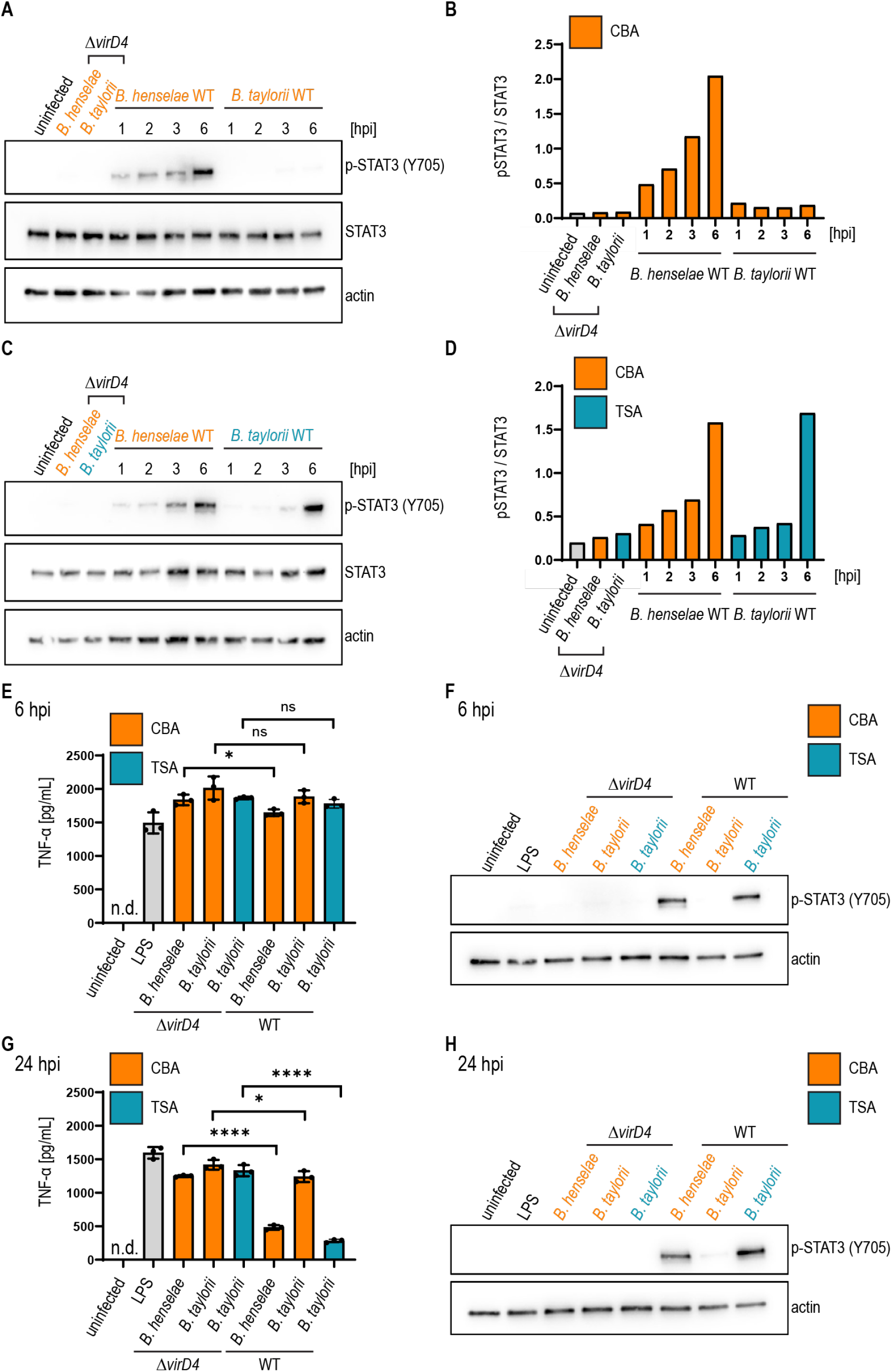
*B. taylorii* grown on TSA trigger a stronger STAT3 activation compared to bacteria grown on CBA. (A) JAWS II cells were infected at MOI 50. At the depicted time points cells were harvested and analyzed by immunoblot with specific antibodies against p-STAT3 (Y705), STAT3 and actin. Phosphorylated STAT3 was quantified over the total amount of STAT3. Shown is an immunoblot of JAWSII cell lysates infected with *B. henselae* wild-type or *ΔvirD4, B. taylorii* wild-type or *ΔvirD4* grown on CBA (orange). Cells were infected for 24 h with *ΔvirD4* mutants. (B) Quantification of immunoblot shown in (A). (C) Shown is an immunoblot of JAWSII cell lysates infected with *B. henselae* wild-type or *ΔvirD4*, *B. taylorii* wild-type or *ΔvirD4* grown on TSA (blue). Cells were infected for 24 h with *ΔvirD4* mutants. (D) Quantification of immunoblot shown in (C). (E) and (G) JAWS II dendritic cells were infected as described in (A). During the last two hours of infection, cells were treated with 100 ng/mL LPS. (E) 6 hpi supernatant was harvested and TNF-α concentration was assessed by ELISA. (F) Cells in (E) were analyzed by Western Blot for phosphorylated STAT3 (Y705). Actin was used as loading control. (G) After 24 hpi TNF-α concentration in the cell culture supernatant was assessed by ELISA. (H) Cells in (G) were analyzed by Western Blot for phosphorylated STAT3 (Y705). Data for immunoblots were acquired by pooling three technical replicates. Data from one representative experiment (n = 3) are presented.

To exclude potential influence on the STAT3 phosphorylation mediated by other effectors present in *B. taylorii*, we infected JAWS II cells with mutants lacking single *bep* genes. *B. taylorii* encodes five Beps, namely BepA-BepI. BepA, BepC and BepI harbor an N-terminal FIC (filamentation induced by cyclic AMP) domain. BepD and BepF contain EPIYA-related motifs in their N-terminus (suppl. Figure S1C). However, BepD_*Bta*_ is the only effector, which induces STAT3 phosphorylation in infected JAWS II cells (suppl. Figure S1D).

Next, we tested if *B. taylorii* downregulates the secretion of pro-inflammatory cytokines in a VirB/VirD4 T4SS-dependent manner as previously published for *B. henselae* (Sorg et al., 2020). We infected JAWS II for 6 h or 24 h with *B. taylorii* wild-type and the *ΔvirD4* mutant. Corresponding *B. henselae* strains served as controls. During the last 2 h of infection, JAWS II cells were co-stimulated with *Escherichia coli* LPS as potent TLR4 ligand to increase TNF-α secretion (Sorg et al., 2020). Cells infected with *B. henselae* wild-type secreted significantly lower TNF-α concentrations compared to the *ΔvirD4* mutant at 6 and 24 hpi. While no discernable inhibitory effect was displayed 6 hpi, *B. taylorii* recovered from CBA plates impaired TNF-α secretion at 24 hpi in a VirD4-dependnet manner. Surprisingly, bacteria recovered from TSA plates did not impair the TNF-α secretion after 6 hpi, although the STAT3 phosphorylation was induced. At 24 hpi, *B. taylorii* cultured on TSA reduced the TNF-α secretion much more efficiently compared to bacteria grown on CBA plates (Figure 1E-H).

*B. henselae* significantly decreases TNF-α secretion at 6 hpi, whereas infection with *B. taylorii* did not reduce the TNF-α concentration in the supernatant of infected JAWS II cells. One explanation might be an earlier onset of STAT3 activation induced by *B. henselae* than by *B. taylorii.* We thus propose that growing *B. taylorii* on TSA plates results in a more efficient downregulation of the innate immune response *in vitro*, while effector translocation into eukaryotic host cells might still occur later compared to infections with *B. henselae*.

### 3.2 Implementing the split-NanoLuc translocation assay to study effector translocation via the VirB/VirD4 T4SS

We aimed at improving the effector translocation via the VirB/VirD4 T4SS of *B. taylorii.* To study effector translocation under various conditions, we implemented the split NanoLuc (NLuc) luciferase-based translocation assay for the VirB/VirD4 T4SS in *Bartonella.* This assay was developed to assess effector injection via the type III secretion system of *Salmonella enterica* serovar Typhimurium in HeLa cells (Westerhausen et al., 2020). Split NLuc is composed of a small fragment (HiBiT, 1.3 kDa), which is fused to the bacterial effectors, and a larger fragment (LgBiT, 18 kDa), which is stably expressed in RAW264.7 macrophages (RAW LgBiT). Different studies already demonstrated successful infection of murine macrophage cell lines by *B. henselae* (Musso et al., 2001; Sorg et al., 2020). Moreover, we already showed that BepD of *B. taylorii* is triggering STAT3 phosphorylation in eukaryotic host cells (suppl. Figure S1F). Therefore, we designed fusion proteins harboring HiBiT and a triple-FLAG epitope tag at the N-terminus of BepD_*Bhe*_ or BepD_*Bta*_ that are expressed under the control of an IPTG-inducible promoter. The fusion proteins were ectopically expressed in the translocation-deficient *ΔvirD4* mutants or the Bep-deficient mutant of *B. henselae (ΔbepA-G)* or *B. taylorii (ΔbepA-I).* As control, bacteria expressing only the HiBiT-FLAG fragment were created. The expression of *pHiBiT-FLAG-bepD_Bhe_* and *pHiBiT-FLAG-bepD_Bta_* after 1 h IPTG-induction was confirmed by immunoblotting using the triple-FLAG epitope tag in bacterial lysates (suppl. Figure S2A and S2B). We could not detect the HiBiT-FLAG fragment (estimated mass 4.3 kDa), most likely due to the low molecular mass and possible degradation. Next, we tested whether the HiBiT-FLAG fragment, HiBiT-FLAG-BepD_*Bhe*_ and HiBiT-FLAG-BepD_*Bta*_ can complement LgBiT to a functional luciferase. We could detect high luminescent signals of lysed bacteria expressing either HiBiT-FLAG-BepD_*Bhe*_ or HiBiT-FLAG-BepD_*Bta*_. Expression of the HiBiT-FLAG fragment also lead to detectable albeit lower luminescent signal (suppl. Figure S2C and S2D).

To estimate the translocation of the HiBiT fused effectors, we infected RAW LgBiT macrophages for 24 h. Complementation of LgBiT with HiBiT to a functional luciferase should only occur if host cells were infected with *Bartonella* strains harboring a functional T4SS. The luciferase converts the substrate furimazine into a bioluminescent signal (Figure 2A). We observed increased luminescent signals inside host cells after infection with *B. henselae ΔbepA-G pHiBiT-FLAG-bepD_Bhe_* (suppl. Figure S2E) or *B. taylorii ΔbepA-I pHiBiT-FLAG-bepD_Bta_* (suppl. Figure S2F), which increased in a MOI-dependent manner. With MOI 50 or higher, significantly increased signals compared to the translocation-deficient mutants were observed for *B. henselae* and *B. taylorii* infection. Therefore, the following experiments were performed at MOI 50.

**Figure 2:**
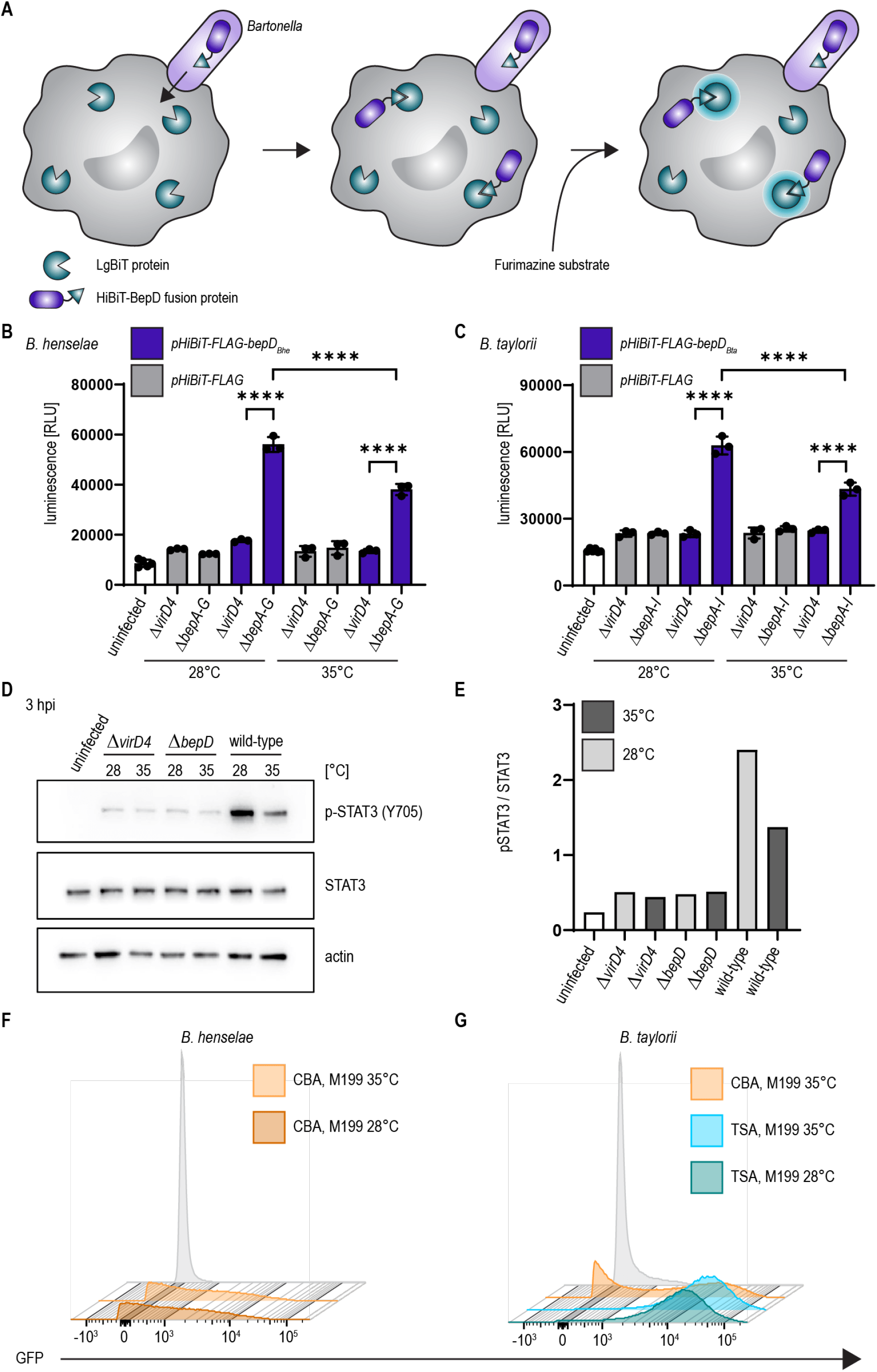
Temperature shift increases effector translocation. (A) Schematic overview showing the split NLuc assay principle. Bacteria were allowed to infect RAW LgBiT macrophages for 24 h. HiBiT-BepD was translocated inside host cells via the VirB/VirD4 T4SS. The substrate Furimazine was added and luminescence measured. (B) Bacteria were cultured in M199 + 10% FCS for 24 h at 28°C or 35°C prior to infection. RAW LgBiT macrophages were infected at MOI 50 for 24 h with *B. henselae ΔbepA-G* or *ΔvirD4* containing *pHiBiT-bepD_Bhe_* (blue) or *pHiBiT* (grey). Luminescence of the complemented split NLuc was measured. (C) RAW LgBiT were infected for 24 h with *B. taylorii ΔbepA-I* or *ΔvirD4* containing *pHiBiT-bepD_Bta_* (blue) or the control *pHiBiT-FLAG* (grey) either cultured in M199 + 10% FCS at 28°C or 35°C for 24 h prior to infection. Luminescence of the complemented split NLuc was measured. (D) RAW 264.7 macrophages were infected for 3 h at MOI 50 with *B. taylorii* wild-type, the BepD-deficient mutant *ΔbepD* or the translocation-deficient mutant *ΔvirD4.* Bacteria were cultured in M199 + 10% FCS at 28°C or 35°C for 24 h prior to infection. Cell lysates were analyzed by Western Blot for phosphorylated STAT3 (Y705), STAT3 and actin. Immunoblot analyzing cell lysates of cells infected for 3 h with bacteria grown at 28°C (light grey) or 35°C (dark grey). (E) Quantification of pSTAT3 signal over STAT3 control of immunoblot shown in (D). Bacteria were grown at 28°C (light grey) or 35°C (dark grey). (F+G) *B. henselae* or *B. taylorii* expressing GFP under the corresponding *virB2 promoter* on a plasmid were grown on CBA (orange) or TSA (blue) plates and cultured for 24 h in M199 + 10% FCS at 28°C or 35°C. GFP expression was analyzed by FACS measurement. Bacteria containing the empty plasmid (pCD366, grey) were used as control. Data for immunoblots were acquired by pooling three technical replicates. All experiments were performed in three independent biological replicates. Data were analyzed using one-way ANOVA with multiple comparisons (Tukey’s multiple comparison test), **** p < 0.0001.

Bartonellae are transmitted via blood-sucking arthropods and experience a temperature shift during their infection cycle from ambient temperature of the arthropod vector to warm-blooded body temperature of the mammalian hosts. *Bartonella* virulence factors have been shown to undergo differential regulation in response to temperature (Abromaitis and Koehler, 2013; Tu et al., 2016). To assess the influence of the temperature on the translocation efficiency, we cultivated the bacteria at 28°C or 35°C in M199 + 10% FCS before infection. RAW LgBiT macrophages were infected at MOI 50 for 24 h. We observed significant higher luminescence compared to the *ΔvirD4* mutants if cells were infected with *B. henselae ΔbepA-G pHiBiT-bepD_Bhe_* (Figure 2B) or *B. taylorii ΔbepA-I pHiBiT-bepD_Bta_* (Figure 2C). The luminescent signals significantly increased if bacteria were cultured at 28°C prior to infection instead of 35°C for both *Bartonella* strains (Figure 2B and 2C).

We also investigated the influence of decreased temperature in *B. taylorii* overnight cultures on the BepD-dependent STAT3 activation in RAW macrophages. While infection with the translocation-deficient *ΔvirD4* mutant or the bacteria lacking *bepD* did not trigger STAT3 activation, we found higher levels of phosphorylated STAT3 at 3 hpi if *B. taylorii* wild-type was cultured at 28°C. The amount of pSTAT3 was quantified over STAT3 (Figure 2D and 2E). A similar phenotype was also observed 6 hpi (suppl. Figure 2G and 2H), although less prominently. This might indicate that BepD translocation is improved at earlier time points if bacteria are cultured at lower temperatures. The incubation at 28°C prior to infection triggers a stronger STAT3 phosphorylation *in vitro*, which seems to correlate with a higher translocation of BepD into host cells.

### 3.3 Increased effector translocation correlates with upregulated expression of the VirB/VirD4 T4SS

Changing the agar plates from CBA to TSA and incubating the overnight culture in M199 at 28°C instead of 35°C markedly improved the luminescent signal in the translocation assay and the STAT3 activation in infected RAW macrophages. Next, we tested whether the enhanced downregulation of the innate immune response after infection with *B. taylorii* recovered from TSA correlates with an upregulation of the VirB/VirD4 T4SS. We generated reporter strains expressing GFP under the control of the *virB2* promoter *(PvirB2)* of *B. henselae* or *B. taylorii*, driving expression of the *virB* operon. The expression of the GFP promoter fusions in *Bartonella* carrying the*pCD366-derived* reporter plasmids was probed using flow cytometry. As shown previously for *B. henselae* (Quebatte et al., 2010; Harms et al., 2017b) only part of the *B. taylorii* population grown on CBA expressed GFP. In comparison, *B. taylorii* cultured on TSA displayed higher fluorescence with almost the entire population being GFP-positive, indicating that these bacteria upregulate expression of the VirB/VirD4 T4SS. However, the incubation at lower temperature did not significantly change the GFP expression (Figure 2F and 2G). The VirB/VirD4 T4SS expression appears to be similar compared to bacteria cultured at 35°C, although we observed increased effector translocation when the bacteria were cultured at lower temperature prior to infection.

Furthermore, we wanted to test, whether the improved culture conditions (Figure 3A) had an impact on the TNF-α secretion after infection by *B. taylorii* at early time points. Thus, we infected RAW macrophages with *B. taylorii* wild-type, the Bep-deficient mutant (*ΔbepA-I)*, a strain lacking only BepD (*ΔbepD*) or the translocation-deficient mutant *ΔvirD4*. The BepD-dependent STAT3 phosphorylation was observed in cells infected with the wild-type at 6 hpi. As expected, the mutants lacking BepD or VirD4 did not trigger increased STAT3 activation (Figure 3B). Compared to wild-type infections, cells infected with the *ΔvirD4* mutant secreted significantly higher levels of TNF-α secretion. Surprisingly, the TNF-α levels of *ΔbepD* and *ΔbepA-I* strains were lowered to similar extend (Figure 3C). 20 h after infection, macrophages infected with the *ΔbepD* and *ΔbepA-I*mutants showed elevated TNF-α secretion compared to wild-type infected cells. However, cells infected with the translocation-deficient *ΔvirD4* mutant secreted significantly higher levels of TNF-α compared to both *Δbep* mutants (Figure 3E), although these strains are unable to trigger STAT3 activation to the same extend as the wild-type (Figure 3D).

**Figure 3:**
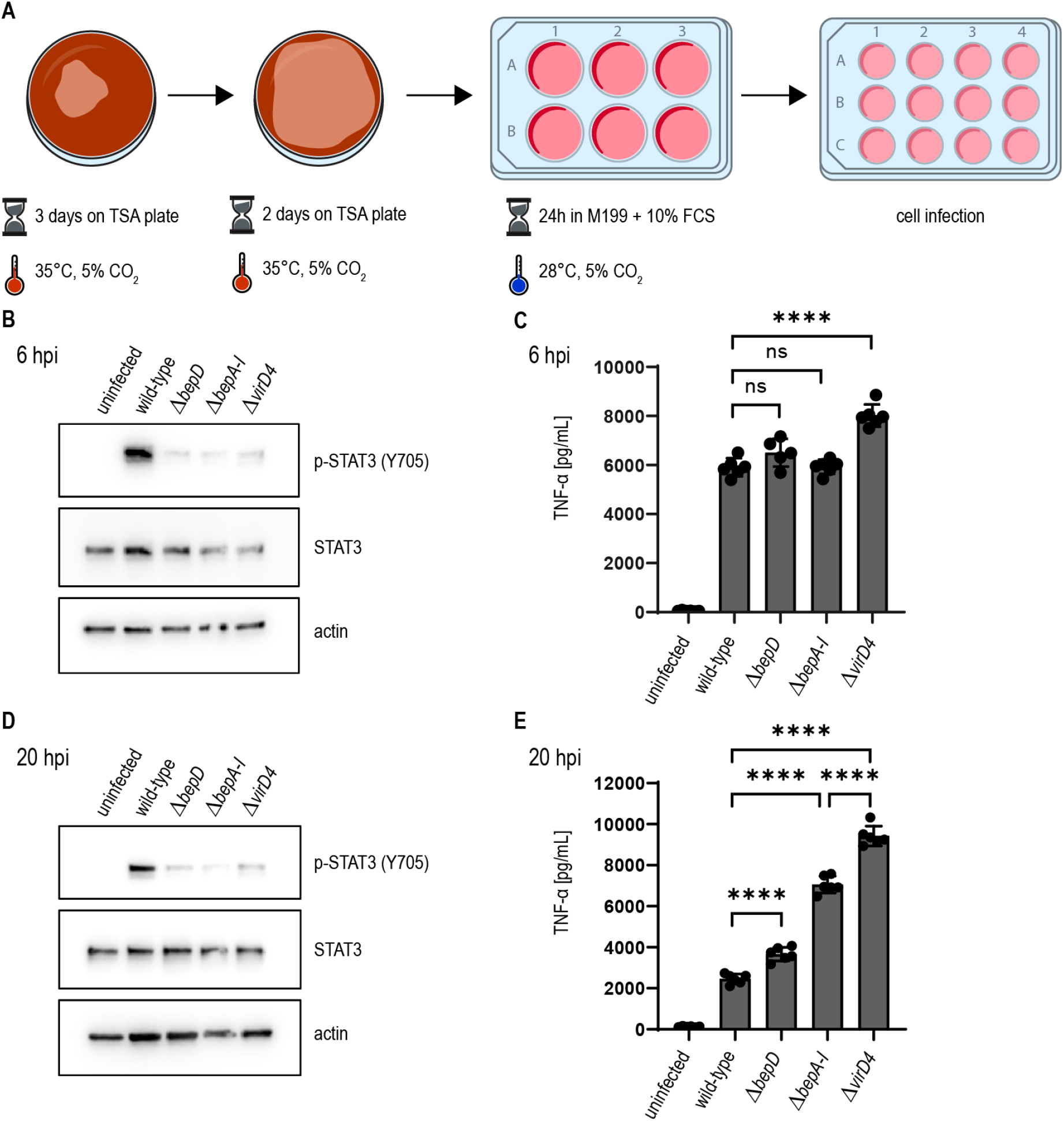
*B. taylorii* efficiently downregulates the innate immune response using the novel *in vitro* infection protocol. (A) Scheme of *B. taylorii* culture conditions used for infection. (B-E) RAW 264.7 macrophages were infected at MOI 50 with *B. taylorii* wild-type, the *ΔbepD* or *ΔbepA-I* mutants or the translocation-deficient mutant *ΔvirD4.* Secreted TNF-α was quantified by ELISA. Cells were harvested, lysed, and analyzed by immunoblot using specific antibodies against p-STAT3 (Y705), STAT3 and actin. (B) Immunoblot of cellular lysates after 6 h infection. (C) TNF-α secreted by cells in (B) was quantified by ELISA. (D) Immunoblot of RAW macrophages infected for 20 h. (E) TNF-α secreted by cells in (D) was quantified by ELISA. Data for immunoblots were acquired by pooling three technical replicates. Data representative for three independent biological replicates. Data were analyzed using one-way ANOVA with multiple comparisons (Tukey’s multiple comparison test), ns = not significant, **** p < 0.0001.

Our data provides evidence that the cultivation of *B. taylorii* prior to infection determines its capacity to dampen the innate immune response *in vitro*. We could correlate this phenotype to an increased expression of the VirB/VirD4 T4SS if bacteria are cultured on TSA plates. However, the lowered temperature seems to not affect the expression of the T4SS indicating that there might be other regulation mechanisms involved, e.g. the effector expression or assembly of the translocation machinery.

### 3.4 Bacterial culture conditions optimized for VirB/VirD4 expression correlate with high infectivity in the mouse model

Finally, we tested whether our adapted growth conditions also influence *B. taylorii in vivo* infections. Previous studies described that mice inoculated with 10^7^ colony forming units (cfu) bacteria remained abacteremic until 5-7 days post infection (dpi). The infection peaked around day 12-14 days with approximately 10^5^ bacteria per ml blood. The bacteremia is cleared within 50 days (Siewert et al., 2021), displaying similar kinetics compared to the infection of rats with *B. tribocorum* (Okujava et al., 2014). We compared the course of bacteremia in wild-type C57BL/6 mice inoculated with different cfus of *B. taylorii* cultured on CBA or TSA plates. Bacteria cultured on TSA plates caused bacteremia independent of the amount of inoculated bacteria, leading to infection rates of 100% even with an inoculum of 10^2^. Bacteria grown on CBA plates showed reduced infectivity (Figure 4A). Interestingly, if the bacteria were able to invade the blood stream, the bacteremia kinetics were similar independent on the growth conditions and inoculum (Figure 4B-4D).

**Figure 4:**
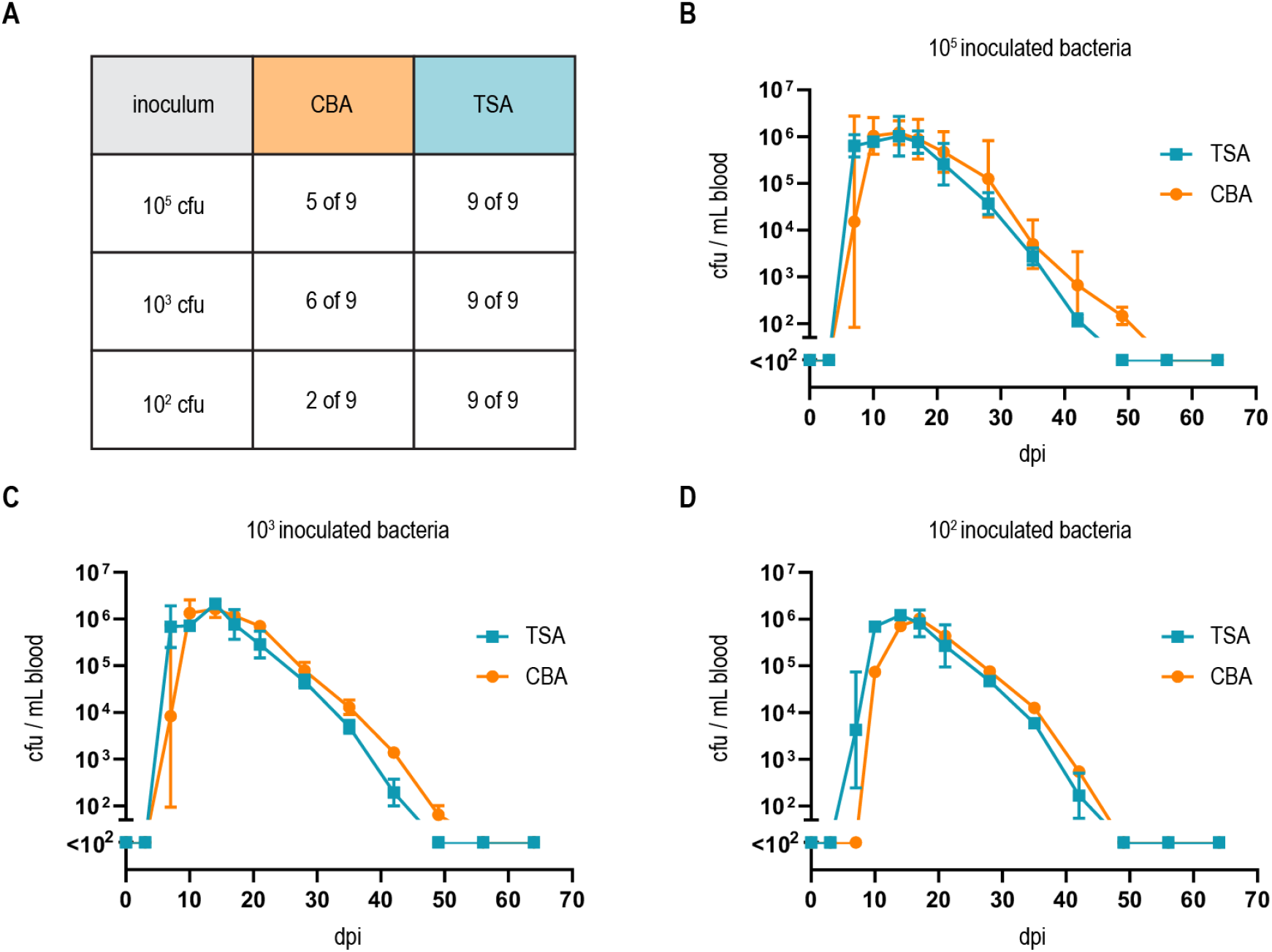
Growth on TSA primes *B. taylorii* for high infectivity in mice. (A) Table represents number of mice developing bacteremia vs mice remaining abacteremic after infection with *B. taylorii* for 9 animals per condition. C57BL/6 mice were infected *i.d.* with 10^2^, 10^3^ or 10^5^ cfu of *B. taylorii* wild-type either grown on CBA (orange) or TSA (blue) plates. Shown are data pooled from three independent experiments. Bacteremia kinetics in infected mice shown for (B) 10^5^, (C) 10^3^ and (D) 10^2^. At several time points after infection, blood was drawn from the tail vein and plated on CBA plates to assess the amount of bacteria inside the blood. Data plotted as mean bacteremia. Figures show representative data from at least three independent experiments.

## 4 Discussion

Typically, *in vitro* studies focusing on the effector translocation via the VirB/VirD4 T4SS of *Bartonella* were conducted with *B. henselae*. In contrast, *in vivo* studies were mostly performed with rodent-specific strains, while *in vitro* infection protocols for those bacteria were missing. Our study characterized the mouse-specific *B. taylorii* strain IBS296 as a suitable model to study effector translocation *in vitro*. The main advantage over previously described species is the use of a murine infection model (Siewert et al., 2021), allowing robust and easy to perform *in vitro* and *in vivo* studies with the same strain.

Growing bacteria on TSA allowed a drastic reduction of the inoculum *in vivo* while the infection rate in C57BL/6 mice remained unchanged. Compared to high-dose infections (10^7^ cfu, (Siewert et al., 2021)), bacteremia kinetics were similar in onset, duration and clearance if mice were infected with 10^2^ cfu. The lower infection dose might better reflect the natural infection via bacteria-containing feces of the arthropod vector scratched into the skin. In experimentally infected fleas the amount of bacteria within their faeces peaked with an average of 9 x 10^4^ bacteria, which drastically decreased several days after infection (Bouhsira et al., 2013). Thus, we consider an infection with 10^2^-10^4^ bacteria as physiological.

We introduced the split NLuc translocation assay as a tool to study effector translocation into host cells via the VirB/VirD4 T4SS. The split NLuc translocation assay had previously been described for studying protein trafficking in mammalian cells (Rouault et al., 2017) and protein secretion in Gram-positive bacteria (Wang et al., 2018). Protein secretion to the periplasm via the Sec system or into host cells via T3SS was shown by utilizing similar approaches in Gram-negative bacteria (Pereira et al., 2019; Westerhausen et al., 2020). Adopting this translocation assay for studying effector translocation via the VirB/VirD4 T4SS in *B. taylorii*, we could show that incubating the bacteria at 28°C instead of 35°C prior to infection improved effector translocation. Additionally, we observed increased STAT3 activation in response to effector translocation when cells were infected with bacteria cultured at lower temperatures. Temperature shift from ambient temperature to 37°C has been shown to regulate virulence factors in several pathogens, such as *Salmonella, Shigella* and *Yersinia* (Cornelis et al., 1987; Prosseda et al., 1998; Shapiro and Cowen, 2012; Lam et al., 2014). In the human-specific pathogen *B. quintana* the alternative sigma factor RpoE is upregulated at lower temperature and high haemin concentrations (Abromaitis and Koehler, 2013). These conditions recapitulate the environment of the arthropod gut and support a role in the adaptation to the vector. In *B. henselae*, RpoE was also found to negatively regulate the transcription of *badA*, which encodes the BadA adhesin (Tu et al., 2016). BadA is an important virulence factor that interacts with endothelial cells in the mammalian host (Riess et al., 2004). In contrast, BadA was described to negatively affect the function of the VirB/VirD4 T4SS (Lu et al., 2013). No evidence was reported that RpoE has direct effects on the *virB* operon as neither the genomic deletion nor ectopic overexpression did influence *virB* promoter activity (Quebatte et al., 2013). However, we cannot exclude that RpoE might affect the function of the VirB/VirD4 T4SS via other regulation mechanisms. We observed upregulated expression of the VirB/VirD4 T4SS in *B. taylorii* cultured on TSA, which might influence the expression of BadA, therefore allowing to monitor effector translocation *in vitro*.

The expression and activation of the VirB/VirD4 T4SS in *B. henselae* is regulated by the alternative sigma factor RpoH1. Levels of RpoH1 are under the control of the stringent response, an adaptive mechanism that allows pathogens to respond to changes in the microenvironments (Dalebroux et al., 2010; Quebatte et al., 2013). Quebatte et al. showed that the stringent response components SpoT and DksA are key regulators of *B. henselae* virulence by controlling the levels of RpoH1 (Quebatte et al., 2013). Additionally, induction of the VirB/VirD4 T4SS also requires an active BatR/BatS two-component system (TCS). The BatR/BatS TCS is activated in neutral pH range (pH 7.0 to 7.8) suggesting that this system is discriminating between the mammalian host (neutral pH in blood and most tissues) and the midgut of the arthropod vector (alkaline pH) (Quebatte et al., 2010). Genes of the BatR regulon and the SpoT/DksA/RpoH1 are conserved amongst Bartonellae (Quebatte et al., 2010; Quebatte et al., 2013), indicating that the expression of the VirB/VirD4 T4SS relies on the same pathways in various *Bartonella* species. However, environmental signals and nutritional states triggering these pathways are not fully understood. We showed that the activation of the VirB/VirD4 T4SS in *B. taylorii* required different culture conditions compared to *B. henselae*. It is thus tempting to speculate that *B. taylorii* may have different metabolic capabilities as *B. henselae* resulting in the observed differences in VirB/VirD4 regulation.

We could show that *B. taylorii* lacking all Beps impaired the TNF-α secretion to a larger extend than the effector translocation-deficient *ΔvirD4* mutant. The TNF-α secretion is likewise impaired following infection with the *ΔbepD* and the full Bep deletion mutant *ΔbepA-I* at 6 hpi, but significantly increased in cells infected with the *ΔvirD4* mutant. Furthermore, we saw differences in the TNF-α secretion when comparing *ΔbepA-I* and *ΔvirD4* mutant at later time points. This observation suggests that some *Bartonella* species harbour at least one additional effector that is translocated via the VirB/VirD4 T4SS. All Beps harbor a bipartite secretion signal composed of one or several Bep intracellular delivery (BID) domains followed by a positively charged tail sequence. It was shown that complete or partial deletion of the BID domain strongly impairs the secretion via the VirB/VirD4 T4SS (Schulein et al., 2005; Schmid et al., 2006b). However, other bacteria expressing a homologous T4SS, like *Agrobacterium tumefaciens*, secrete effector proteins without BID domains. Positively charged amino acids in the C-terminus of recognized effectors serve as translocation signal. For example, the last C-terminal 20 amino acids of VirF of *A. tumefaciens* are sufficient to serve as translocation signal (Vergunst et al., 2005). Here we provide data suggesting that *B. taylorii* might harbour other effector proteins probably lacking BID domains, which might be translocated by the VirB/VirD4 T4SS. We suggest to employ the NLuc translocation assay to study the translocation of putative new effectors in *Bartonella* as this system provides high signal-to noise ratios, is easy to perform and allows high-throughput screening.

Taken together, this study characterized *B. taylorii* IBS296 as suitable model organism allowing for the first time to directly compare host-pathogen interaction *in vitro* and in rodent hosts using the same *Bartonella* species.

## 6 Author Contributions

K. F. and C. D. conceptualized the studies and designed all experiments. K. F. designed the figures. K. F. (cell infections, Nano-Luc translocation assay, animal work, flow cytometry, cloning, western blotting, ELISA); A. B. (cell infections, western blotting, ELISA); M. O. (Nano-Luc translocation assay, cloning); A. W. (cloning); E. B. (generation LgBiT RAW macrophages); S. M. (cloning); S. W. (experimental design) conducted the experiments, collected and analysed the data. K. F. and C. D. wrote the manuscript. All authors have read and approved the final version of the manuscript.

## 8 Acknowledgements

We specially want to thank Lena Siewert, Jaroslaw Sedzicki and Maxime Québatte for helpful comments and critical reading of the manuscript.

---

**Figure S1:**
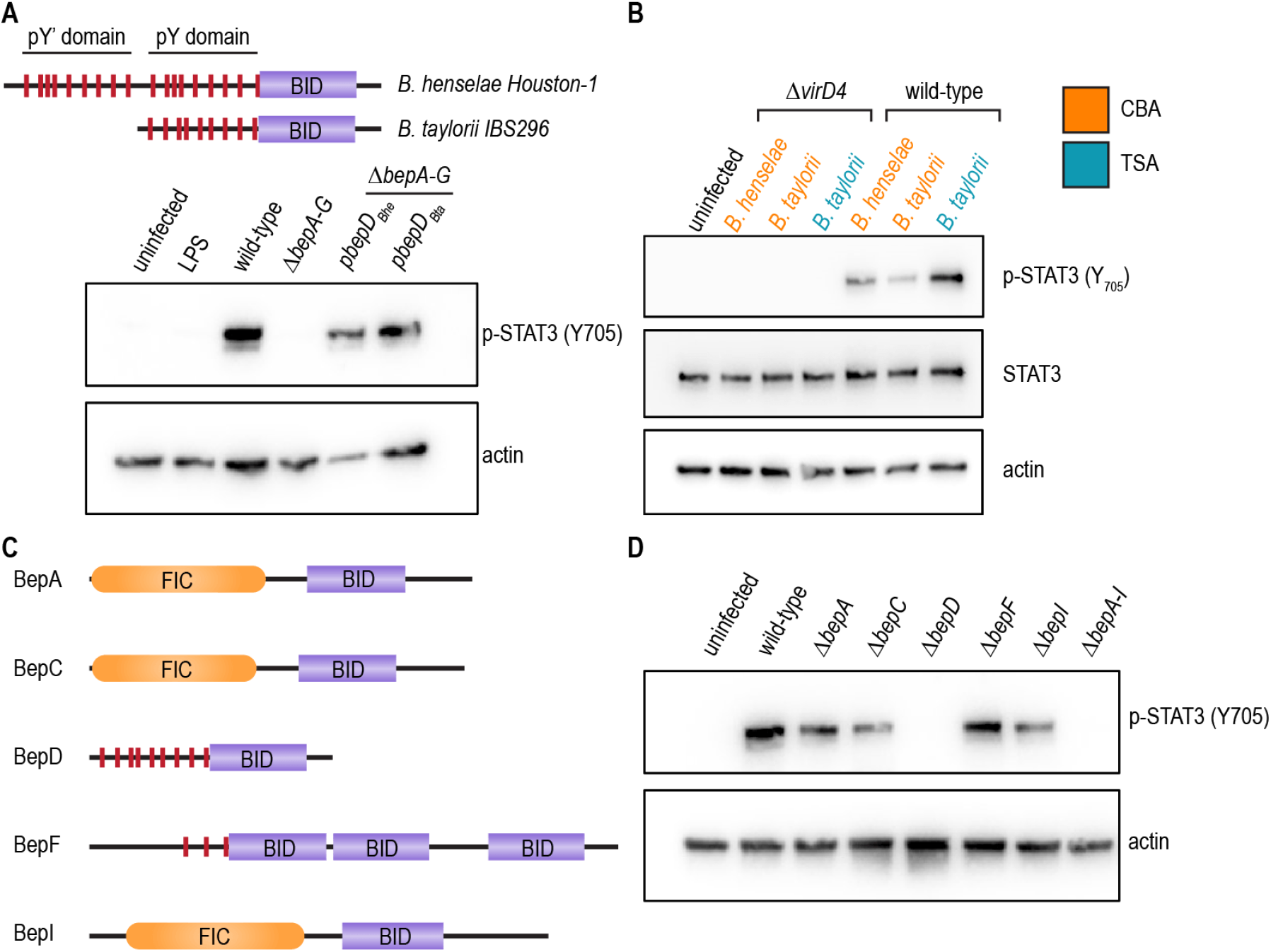
BepD_*Bta*_ ortholog of *B. taylorii* activates STAT3. (A) BepD domain architecture of *B. henselae* Houston-1 and *B. taylorii* IBS296. *B. henselae* harbors 18 tyrosine residues (red) embedded within the two pY’ and pY domains. BepD of *B. taylorii* contains only the pY domain with 9 tyrosine residues. JAWS II dendritic cells were infected at MOI 50 with *B. henselae* wild-type, the Bep-deficient strain *ΔbepA-*G, its BepD_*Bhe*_-expressing derivative *ΔbepA-G pbepD_Bhe_* or its BepD_*Bta*_-expressing derivative *ΔbepA-G pbepD_Bta_*. At 6 hpi, cells were harvested, lysed, and analyzed by immunoblot with specific antibodies against phosphorylated STAT3 (Y705) and actin. (B) Data contributes to figure 1C-F. JAWS II cells were infected at MOI 50 for 24 h with the wild-type or the *ΔvirD4* mutant of *B. henselae* or *B. taylorii* grown on CBA (orange) or TSA (blue). Cells were harvested, lysed and analyzed by immunoblot using specific antibodies against p-STAT3 (Y705), STAT3 and actin. Total amount of STAT3 was used to quantify the phosphorylation of STAT3. (C) Domain architecture of the Bep repertoire present in *B. taylorii*. FIC domains are displayed in orange, BID domains shown in purple and phosphorylation motifs are shown as red, vertical lines. (D) JAWS II dendritic cells were infected at MOI 50 with *B. taylorii* wild-type, the Bep-deficient strain *ΔbepA-I* or single-*bep* deletions. At 6 hpi cells were harvested, lysed and analyzed by Western Blot for phosphorylated STAT3 (Y705) and actin. Data was acquired by pooling three technical replicates and performed in three independent biological experiments. FIC = filamentation induced by cyclic AMP; BID = *Bartonella* effector protein intracellular delivery

**Figure S2:**
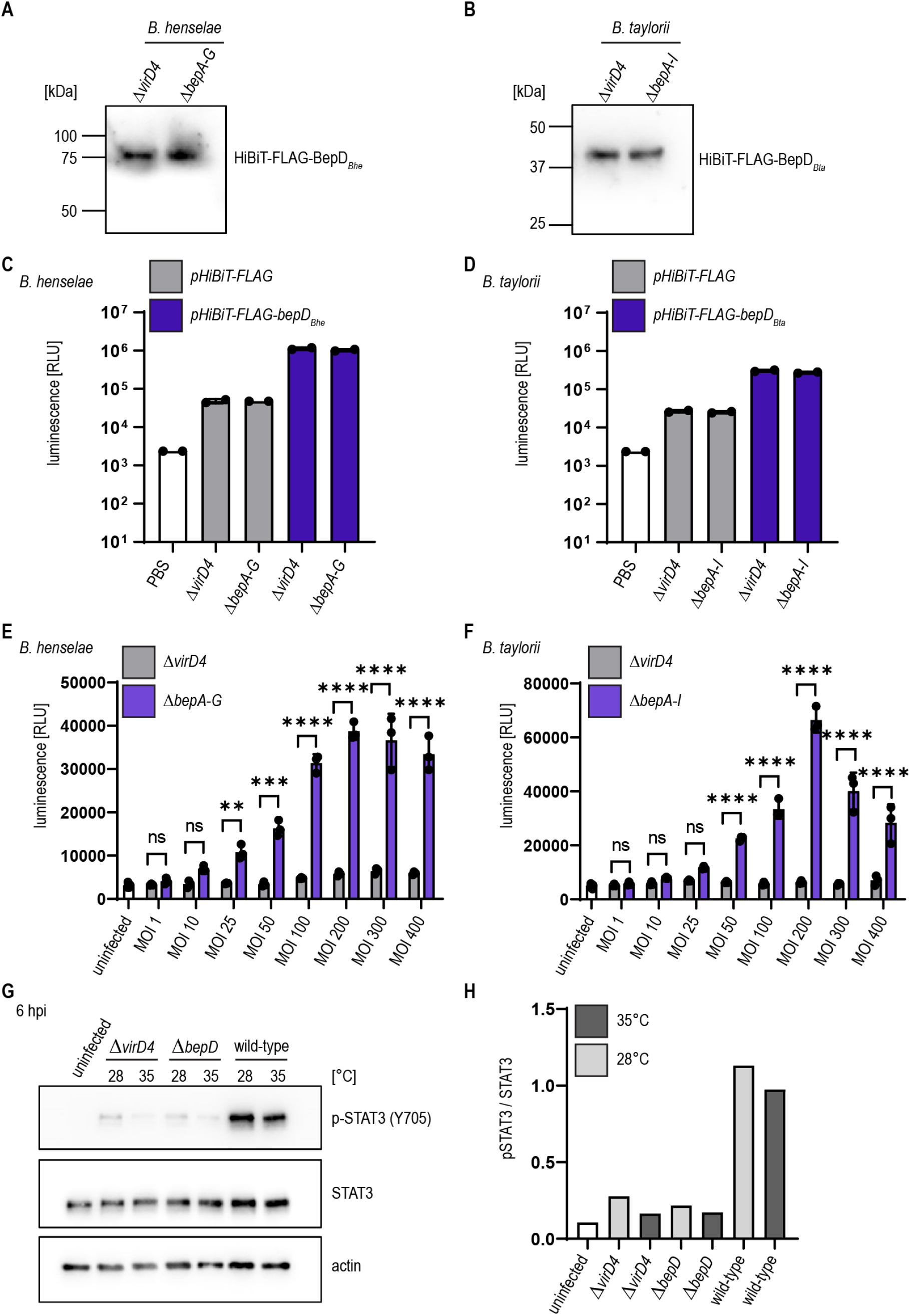
BepD_*Bhe*_ and BepD_*Bta*_ are translocated inside RAW macrophages in a *virD4*- and dose-dependent manner. Immunoblot using specific antibody against the FLAG-epitope. The calculated molecular mass of (A) HiBiT-FLAG-BepD_*Bhe*_ is 63.6 kDa and the calculated molecular mass of (B) HiBiT-FLAG-BepD_*Bta*_ is 42.6 kDa. The interaction of HiBiT-FLAG (grey), HiBiT-FLAG-BepD_*Bhe*_ or HiBiT-FLAG-BepD_*Bta*_ (both shown in blue) with LgBiT was tested using the Nano-Glo HiBiT lytic detection system. Lysed bacteria were supplemented with the purified LgBiT protein and the substrate and luminescence measured using the Synergy H4 plate reader for (C) *B. henselae* or (D) *B. taylorii.* (E) RAW LgBiT macrophages were infected with with *B. henselae ΔbepA-G* (blue) or *ΔvirD4* (grey) containing *pHiBiT-FLAG-bepD_Bhe_* for 24 h with the indicated MOIs, washed and supplemented with the NLuc substrate. Luminescence was measured in the Synergy H4 plate reader. (F) RAW LgBiT macrophages were infected with *B. taylorii ΔbepA-I* (blue) or *ΔvirD4* (grey) containing *pHiBiT-FLAG-bepD_Bta_* and luminescence measured after 24 hpi. (G) RAW 264.7 macrophages were infected at MOI 50 with *B. taylorii* wild-type, the BepD-deficient mutant *ΔbepD* or the translocation-deficient mutant *ΔvirD4.* Cell lysates were analyzed by Western Blot for phosphorylated STAT3 (Y705), STAT3 and actin. Immunoblot analyzing cell lysates of cells infected for 6 h with bacteria grown at 28°C (light grey) or 35°C (dark grey). (E) Quantification of pSTAT3 signal over STAT3 control of immunoblot shown in (D). Data was analyzed using one-way ANOVA with multiple comparisons (Tukey’s multiple comparison test), ns = not significant, ** p < 0.01, *** p < 0.001, **** p < 0.0001

## References

Abromaitis, S., and Koehler, J.E. (2013). The Bartonella quintana extracytoplasmic function sigma factor RpoE has a role in bacterial adaptation to the arthropod vector environment. J Bacteriol 195(11), 2662–2674. doi: 10.1128/JB.01972-12.

Berge, C., Waksman, G., and Terradot, L. (2017). Structural and Molecular Biology of Type IV Secretion Systems. Curr Top Microbiol Immunol 413, 31–60. doi: 10.1007/978-3-319-75241-9_2.

Bouhsira, E., Franc, M., Boulouis, H.J., Jacquiet, P., Raymond-Letron, I., and Lienard, E. (2013). Assessment of persistence of Bartonella henselae in Ctenocephalides felis. Appl Environ Microbiol 79(23), 7439–7444. doi: 10.1128/AEM.02598-13.

Boulouis, H.J., Barrat, F., Bermond, D., Bernex, F., Thibault, D., Heller, R., et al. (2001). Kinetics of Bartonella birtlesii infection in experimentally infected mice and pathogenic effect on reproductive functions. Infect Immun 69(9), 5313–5317. doi: 10.1128/iai.69.9.5313-5317.2001.

Chomel, B.B., Boulouis, H.J., Breitschwerdt, E.B., Kasten, R.W., Vayssier-Taussat, M., Birtles, R.J., et al. (2009). Ecological fitness and strategies of adaptation of Bartonella species to their hosts and vectors. Vet Res 40(2), 29. doi: 10.1051/vetres/2009011.

Chomel, B.B., and Kasten, R.W. (2010). Bartonellosis, an increasingly recognized zoonosis. J Appl Microbiol 109(3), 743–750. doi: 10.1111/j.1365-2672.2010.04679.x.

Chomel, B.B., Kasten, R.W., Floyd-Hawkins, K., Chi, B., Yamamoto, K., Roberts-Wilson, J., et al. (1996). Experimental transmission of Bartonella henselae by the cat flea. J Clin Microbiol 34(8), 1952–1956. doi: 10.1128/JCM.34.8.1952-1956.1996.

Cornelis, G., Vanootegem, J.C., and Sluiters, C. (1987). Transcription of the yop regulon from Y. enterocolitica requires trans acting pYV and chromosomal genes. Microb Pathog 2(5), 367–379. doi: 10.1016/0882-4010(87)90078-7.

Dalebroux, Z.D., Svensson, S.L., Gaynor, E.C., and Swanson, M.S. (2010). ppGpp conjures bacterial virulence. Microbiol Mol Biol Rev 74(2), 171–199. doi: 10.1128/MMBR.00046-09.

Dehio, M., Knorre, A., Lanz, C., and Dehio, C. (1998). Construction of versatile high-level expression vectors for Bartonella henselae and the use of green fluorescent protein as a new expression marker. Gene 215(2), 223–229. doi: 10.1016/s0378-1119(98)00319-9.

Deng, H., Pang, Q., Xia, H., Le Rhun, D., Le Naour, E., Yang, C., et al. (2016). Identification and functional analysis of invasion associated locus B (IalB) in Bartonella species. Microb Pathog 98, 171–177. doi: 10.1016/j.micpath.2016.05.007.

Foil, L., Andress, E., Freeland, R.L., Roy, A.F., Rutledge, R., Triche, P.C., et al. (1998). Experimental infection of domestic cats with Bartonella henselae by inoculation of Ctenocephalides felis (Siphonaptera: Pulicidae) feces. J Med Entomol 35(5), 625–628. doi: 10.1093/jmedent/35.5.625.

Fromm, K., and Dehio, C. (2021). The Impact of Bartonella VirB/VirD4 Type IV Secretion System Effectors on Eukaryotic Host Cells. Frontiers in Microbiology 12. doi: 10.3389/fmicb.2021.762582.

Harms, A., and Dehio, C. (2012). Intruders below the radar: molecular pathogenesis of Bartonella spp. Clin Microbiol Rev 25(1), 42–78. doi: 10.1128/CMR.05009-11.

Harms, A., Liesch, M., Korner, J., Quebatte, M., Engel, P., and Dehio, C. (2017a). A bacterial toxin-antitoxin module is the origin of inter-bacterial and inter-kingdom effectors of Bartonella. PLoS Genet 13(10), e1007077. doi: 10.1371/journal.pgen.1007077.

Harms, A., Segers, F.H., Quebatte, M., Mistl, C., Manfredi, P., Korner, J., et al. (2017b). Evolutionary Dynamics of Pathoadaptation Revealed by Three Independent Acquisitions of the VirB/D4 Type IV Secretion System in Bartonella. Genome Biol Evol 9(3), 761–776. doi: 10.1093/gbe/evx042.

Huarcaya, E., Maguina, C., Merello, J., Cok, J., Birtles, R., Infante, B., et al. (2002). A prospective study of Cat-Scratch Disease in Lima-Peru. Rev Inst Med Trop Sao Paulo 44(6), 325–330. doi: 10.1590/s0036-46652002000600006.

Jiang, X., Shen, C., Rey-Ladino, J., Yu, H., and Brunham, R.C. (2008). Characterization of murine dendritic cell line JAWS II and primary bone marrow-derived dendritic cells in Chlamydia muridarum antigen presentation and induction of protective immunity. Infect Immun 76(6), 2392–2401. doi: 10.1128/IAI.01584-07.

Khalfe, N., and Lin, D. (2022). Diagnosis and interpretation of testing for cat scratch disease. Proc (Bayl Univ Med Cent) 35(1), 68–69. doi: 10.1080/08998280.2021.1984791.

Koesling, J., Aebischer, T., Falch, C., Schulein, R., and Dehio, C. (2001). Cutting edge: antibody-mediated cessation of hemotropic infection by the intraerythrocytic mouse pathogen Bartonella grahamii. J Immunol 167(1), 11–14. doi: 10.4049/jimmunol.167.1.11.

Kunz, S., Oberle, K., Sander, A., Bogdan, C., and Schleicher, U. (2008). Lymphadenopathy in a novel mouse model of Bartonella-induced cat scratch disease results from lymphocyte immigration and proliferation and is regulated by interferon-alpha/beta. Am J Pathol 172(4), 1005–1018. doi: 10.2353/ajpath.2008.070591.

Lam, O., Wheeler, J., and Tang, C.M. (2014). Thermal control of virulence factors in bacteria: a hot topic. Virulence 5(8), 852–862. doi: 10.4161/21505594.2014.970949.

Li, D.-M., Hou, Y., Song, X.-P., Fu, Y.-Q., Li, G.-C., Li, M., et al. (2015). High Prevalence and Genetic Heterogeneity of Rodent-Borne Bartonella Species on Heixiazi Island, China. Applied and Environmental Microbiology 81(23), 7981–7992. doi: doi:10.1128/AEM.02041-15.

Lu, Y.Y., Franz, B., Truttmann, M.C., Riess, T., Gay-Fraret, J., Faustmann, M., et al. (2013). Bartonella henselae trimeric autotransporter adhesin BadA expression interferes with effector translocation by the VirB/D4 type IV secretion system. Cell Microbiol 15(5), 759–778. doi: 10.1111/cmi.12070.

Ma, K.W., and Ma, W. (2016). YopJ Family Effectors Promote Bacterial Infection through a Unique Acetyltransferase Activity. Microbiol Mol Biol Rev 80(4), 1011–1027. doi: 10.1128/MMBR.00032-16.

Mada, P.K., Zulfiqar, H., and Joel Chandranesan, A.S. (2022). “Bartonellosis,” in StatPearls. (Treasure Island (FL)).

Maguina, C., Guerra, H., and Ventosilla, P. (2009). Bartonellosis. Clin Dermatol 27(3), 271–280. doi: 10.1016/j.clindermatol.2008.10.006.

Marlaire, S., and Dehio, C. (2021). Bartonella effector protein C mediates actin stress fiber formation via recruitment of GEF-H1 to the plasma membrane. PLoS Pathog 17(1), e1008548. doi: 10.1371/journal.ppat.1008548.

McCord, A.M., Burgess, A.W., Whaley, M.J., and Anderson, B.E. (2005). Interaction of Bartonella henselae with endothelial cells promotes monocyte/macrophage chemoattractant protein 1 gene expression and protein production and triggers monocyte migration. Infect Immun 73(9), 5735–5742. doi: 10.1128/IAI.73.9.5735-5742.2005.

Musso, T., Badolato, R., Ravarino, D., Stornello, S., Panzanelli, P., Merlino, C., et al. (2001). Interaction of Bartonella henselae with the murine macrophage cell line J774: infection and proinflammatory response. Infect Immun 69(10), 5974–5980. doi: 10.1128/IAI.69.10.5974-5980.2001.

Okujava, R., Guye, P., Lu, Y.Y., Mistl, C., Polus, F., Vayssier-Taussat, M., et al. (2014). A translocated effector required for Bartonella dissemination from derma to blood safeguards migratory host cells from damage by co-translocated effectors. PLoS Pathog 10(6), e1004187. doi: 10.1371/journal.ppat.1004187.

Pereira, G.C., Allen, W.J., Watkins, D.W., Buddrus, L., Noone, D., Liu, X., et al. (2019). A High-Resolution Luminescent Assay for Rapid and Continuous Monitoring of Protein Translocation across Biological Membranes. J Mol Biol 431(8), 1689–1699. doi: 10.1016/j.jmb.2019.03.007.

Prosseda, G., Fradiani, P.A., Di Lorenzo, M., Falconi, M., Micheli, G., Casalino, M., et al. (1998). A role for H-NS in the regulation of the virF gene of Shigella and enteroinvasive Escherichia coli. Res Microbiol 149(1), 15–25. doi: 10.1016/s0923-2508(97)83619-4.

Quebatte, M., Dehio, M., Tropel, D., Basler, A., Toller, I., Raddatz, G., et al. (2010). The BatR/BatS two-component regulatory system controls the adaptive response of Bartonella henselae during human endothelial cell infection. J Bacteriol 192(13), 3352–3367. doi: 10.1128/JB.01676-09.

Quebatte, M., Dick, M.S., Kaever, V., Schmidt, A., and Dehio, C. (2013). Dual input control: activation of the Bartonella henselae VirB/D4 type IV secretion system by the stringent sigma factor RpoH1 and the BatR/BatS two-component system. Mol Microbiol 90(4), 756–775. doi: 10.1111/mmi.12396.

Raschke, W.C., Baird, S., Ralph, P., and Nakoinz, I. (1978). Functional macrophage cell lines transformed by Abelson leukemia virus. Cell 15(1), 261–267. doi: 10.1016/0092-8674(78)90101-0.

Regnath, T., Mielke, M.E., Arvand, M., and Hahn, H. (1998). Murine model of Bartonella henselae infection in the immunocompetent host. Infect Immun 66(11), 5534–5536. doi: 10.1128/IAI.66.11.5534-5536.1998.

Riess, T., Andersson, S.G., Lupas, A., Schaller, M., Schafer, A., Kyme, P., et al. (2004). Bartonella adhesin a mediates a proangiogenic host cell response. J Exp Med 200(10), 1267–1278. doi: 10.1084/jem.20040500.

Rouault, A.A.J., Lee, A.A., and Sebag, J.A. (2017). Regions of MRAP2 required for the inhibition of orexin and prokineticin receptor signaling. Biochim Biophys Acta Mol Cell Res 1864(12), 2322–2329. doi: 10.1016/j.bbamcr.2017.09.008.

Schmid, D., Dengjel, J., Schoor, O., Stevanovic, S., and Munz, C. (2006a). Autophagy in innate and adaptive immunity against intracellular pathogens. J Mol Med (Berl) 84(3), 194–202. doi: 10.1007/s00109-005-0014-4.

Schmid, M.C., Scheidegger, F., Dehio, M., Balmelle-Devaux, N., Schulein, R., Guye, P., et al. (2006b). A translocated bacterial protein protects vascular endothelial cells from apoptosis. PLoS Pathog 2(11), e115. doi: 10.1371/journal.ppat.0020115.

Schmid, M.C., Schulein, R., Dehio, M., Denecker, G., Carena, I., and Dehio, C. (2004). The VirB type IV secretion system of Bartonella henselae mediates invasion, proinflammatory activation and antiapoptotic protection of endothelial cells. Mol Microbiol 52(1), 81–92. doi: 10.1111/j.1365-2958.2003.03964.x.

Schulein, R., and Dehio, C. (2002). The VirB/VirD4 type IV secretion system of Bartonella is essential for establishing intraerythrocytic infection. Mol Microbiol 46(4), 1053–1067.

Schulein, R., Guye, P., Rhomberg, T.A., Schmid, M.C., Schroder, G., Vergunst, A.C., et al. (2005). A bipartite signal mediates the transfer of type IV secretion substrates of Bartonella henselae into human cells. Proc Natl Acad Sci U S A 102(3), 856–861. doi: 10.1073/pnas.0406796102.

Seubert, A., Schulein, R., and Dehio, C. (2002). Bacterial persistence within erythrocytes: a unique pathogenic strategy of Bartonella spp. Int J Med Microbiol 291(6-7), 555–560. doi: 10.1078/1438-4221-00167.

Shapiro, R.S., and Cowen, L.E. (2012). Thermal control of microbial development and virulence: molecular mechanisms of microbial temperature sensing. mBio 3(5). doi: 10.1128/mBio.00238-12.

Siewert, L.K., Korotaev, A., Sedzicki, J., Fromm, K., Pinschewer, D.D., and Dehio, C. (2021). The <em>Bartonella</em> autotransporter CFA is a protective antigen and hypervariable target of neutralizing antibodies blocking erythrocyte infection. bioRxiv, 2021.2009.2029.462357. doi: 10.1101/2021.09.29.462357.

Sorg, I., Schmutz, C., Lu, Y.Y., Fromm, K., Siewert, L.K., Bogli, A., et al. (2020). A Bartonella Effector Acts as Signaling Hub for Intrinsic STAT3 Activation to Trigger Anti-inflammatory Responses. Cell Host Microbe 27(3), 476–485 e477. doi: 10.1016/j.chom.2020.01.015.

Stepanic, M., Duvnjak, S., Reil, I., Spicic, S., Kompes, G., and Beck, R. (2019). First isolation and genotyping of Bartonella henselae from a cat living with a patient with cat scratch disease in Southeast Europe. BMC Infect Dis 19(1), 299. doi: 10.1186/s12879-019-3929-z.

Tu, N., Lima, A., Bandeali, Z., and Anderson, B. (2016). Characterization of the general stress response in Bartonella henselae. Microb Pathog 92, 1–10. doi: 10.1016/j.micpath.2015.12.010.

Vayssier-Taussat, M., Le Rhun, D., Deng, H.K., Biville, F., Cescau, S., Danchin, A., et al. (2010). The Trw type IV secretion system of Bartonella mediates host-specific adhesion to erythrocytes. PLoS Pathog 6(6), e1000946. doi: 10.1371/journal.ppat.1000946.

Vergunst, A.C., van Lier, M.C., den Dulk-Ras, A., Stuve, T.A., Ouwehand, A., and Hooykaas, P.J. (2005). Positive charge is an important feature of the C-terminal transport signal of the VirB/D4-translocated proteins of Agrobacterium. Proc Natl Acad Sci U S A 102(3), 832–837. doi: 10.1073/pnas.0406241102.

Wagner, A., and Dehio, C. (2019). Role of distinct Type-IV-secretion systems and secreted effector sets in host adaptation by pathogenic Bartonella species. Cell Microbiol, e13004. doi: 10.1111/cmi.13004.

Wagner, A., Tittes, C., and Dehio, C. (2019). Versatility of the BID Domain: Conserved Function as Type-IV-Secretion-Signal and Secondarily Evolved Effector Functions Within Bartonella-Infected Host Cells. Front Microbiol 10, 921. doi: 10.3389/fmicb.2019.00921.

Waksman, G. (2019). From conjugation to T4S systems in Gram-negative bacteria: a mechanistic biology perspective. EMBO Rep 20(2). doi: 10.15252/embr.201847012.

Wang, C., Fu, J., Wang, M., Cai, Y., Hua, X., Du, Y., et al. (2019). Bartonella quintana type IV secretion effector BepE-induced selective autophagy by conjugation with K63 polyubiquitin chain. Cell Microbiol 21(4), e12984. doi: 10.1111/cmi.12984.

Wang, C.Y., Patel, N., Wholey, W.Y., and Dawid, S. (2018). ABC transporter content diversity in Streptococcus pneumoniae impacts competence regulation and bacteriocin production. Proc Natl Acad Sci U S A 115(25), E5776–E5785. doi: 10.1073/pnas.1804668115.

Westerhausen, S., Nowak, M., Torres-Vargas, C.E., Bilitewski, U., Bohn, E., Grin, I., et al. (2020). A NanoLuc luciferase-based assay enabling the real-time analysis of protein secretion and injection by bacterial type III secretion systems. Mol Microbiol 113(6), 1240–1254. doi: 10.1111/mmi.14490.

